# Stochastic optimal control simulations of walking: potential and perspective

**DOI:** 10.64898/2026.03.19.712839

**Authors:** Lars D’Hondt, Maarten Afschrift, Friedl De Groote

## Abstract

Human walking is intrinsically variable. For example, there is considerable stride to stride variability even when walking speed is constant. This variability is due to uncertainty in the sensorimotor system and the environment, and is shaped by both musculoskeletal dynamics (e.g. joint stiffness and damping originating from muscles) and the control strategy used to mitigate the effects of uncertainty. Yet, insight into how sensorimotor noise shapes walking variability is limited due to a lack of experimental methods to assess sensorimotor noise and control strategies during walking. Simulations that account for uncertainty can elucidate how sensorimotor noise affects movement variability but due to numerical challenges, accounting for sensorimotor noise is not common in simulations of walking. Existing simulations have hugely simplified musculoskeletal dynamics (e.g. no muscles), the control policy (e.g. pre-defined feedback loops), or sensorimotor noise sources (e.g. only motor noise). Here, we performed stochastic optimal control simulations of walking based on a model with 9 degrees of freedom and 18 muscles to study how the level of sensory and motor noise influences walking. We solved for feedforward muscle excitations and full-state time-varying feedback gains that minimised expected effort while generating periodic, and hence stable, gait patterns. To enable these simulations, we approximated the state distribution with a Gaussian and used an unscented transform to propagate the state covariance. Resulting optimisation problems were solved with direct collocation. Sensorimotor noise level had a small effect on the mean kinematics but shaped kinematic and muscle activity variability as well as expected effort. Although simulations underestimated the magnitude of experimental positional variability, they captured its structure. In agreement with experimental results, the control policy prioritised limiting variability of centre of mass kinematics and minimal swing foot clearance over limiting joint angle variability. Hence, our simulations suggest that effort minimisation underlies these observations.

**Author summary:** When performing a movement multiple times, each repetition will be slightly different due to random disturbances in the neural signals used to control movement, i.e. sensorimotor noise. Because it is difficult to measure inside the nervous system of a moving person, computer simulations are used to study movement control. They found that both sensorimotor noise and musculoskeletal mechanics determine how people control arm movements and standing. However, there are no simulations of walking that systematically evaluated how sensorimotor noise level influences walking kinematics because they pose computational challenges. Here, we proposed and used an approach for minimal effort simulations of walking in the presence of uncertainty. We imposed forward speed and stability but not kinematics. We found that the level of sensorimotor noise had little effect on the mean movement but a strong effect on the variability and the expected effort. The control strategy prioritised reducing the variability of the centre of mass position and swing foot clearance over reducing the variability of individual joint angles, which is also observed in experiments. Interestingly, strict control of centre of mass position and foot clearance in our simulations emerged from minimising effort.

## Introduction

Human movement is intrinsically variable. For example, walking kinematics varies from stride to stride even when walking at a constant speed. This variability is due to uncertainty in both the sensorimotor system and the environment (1), and is shaped by the control strategy used to mitigate the effects of uncertainty. Yet, our understanding of how uncertainty and control shape walking variability is limited because experimental methods to assess sensorimotor noise and control strategies are limited. Model-based simulations have the potential to elicit causal relationships between uncertainty and movement variability. Yet, there are no simulations of walking that systematically evaluated how sensorimotor noise level influences walking. The high computational cost of simulation methods that account for uncertainty restricts the use of neuromusculoskeletal models that are sufficiently complex to realistically capture walking. Therefore, we present a novel approach to simulate walking in the presence of uncertainty. We used this approach to investigate how sensorimotor noise shapes walking.

It has been demonstrated that accounting for uncertainty is crucial to simulate movement variability and sensorimotor feedback in a range of non-walking movements. Such simulations solve for control policies that minimise a cost in the presence of noise. The cost is typically quantified as the expected task error (e.g. distance from target position in a reaching task), the expected effort, or a combination of both. In feedforward-controlled simulations of fast arm or eye movements, accounting for motor noise was crucial to obtain movement patterns that resembled experimentally observed movement patterns (2,3). When moving slowly, humans use continuous sensory feedback to correct deviations due to noise. Optimal control simulations in the presence of noise yielded feedback policies that allowed variability to accumulate in the task-irrelevant dimensions, rather than to eliminate all variability, in agreement with experimental data for various arm reaching tasks (4). Simulation studies mainly focused on arm movements (2–8) or standing postural control (8–10), and commonly represented the musculoskeletal system by a conceptual model, often with linear dynamics (2–7,9). Computational challenges have limited the use of simulations based on more detailed models with nonlinear dynamics.

Musculoskeletal dynamics influence how uncertainty affects movement. Joint impedance, influenced by muscle co-contraction, helps absorb and dissipate perturbation energy (11,12). While neural feedback acts with a delay, mechanical responses are instantaneous (13,14). Recent optimal control simulations of nonlinear dynamics in the presence of uncertainty, enabled through computational advances, have demonstrated that muscle mechanics influence the optimal control strategy used to stabilise the movement against uncertainty (8). Using a muscle-driven multi-segment arm model yielded more realistic arm reaching trajectories than using a point mass model and these trajectories were influenced by muscle geometry and muscle contractile properties (8). By modelling antagonistic muscles with activation-dependent stiffness, it could be demonstrated that muscle co-contraction – a strategy thought to be energetically costly – can be a minimal effort strategy to stabilise standing postural control (8,10). However, there are no walking simulations that describe musculoskeletal dynamics, neural control, and sensorimotor noise in sufficient detail to gain insights into their interaction during walking.

Simulations of walking have neglected or drastically simplified uncertainty. A first class of simulations does not explicitly model the control policy. Instead, they solve for control parameters that result in a periodic motion with a desired forward velocity, while optimising a performance criterion in the absence of uncertainty (15,16). The resulting optimal control problems can be solved efficiently using direct collocation enabling the use of detailed musculoskeletal models. Such simulations have advanced insight into how musculoskeletal properties and impairments affect the gait pattern (17–20). However, this approach generates only a nominal gait cycle. Hence, it cannot be used to study how uncertainty shapes movement variability. Another class of simulations explicitly models the control policy but typically oversimplifies the control policy and/or does not account for sensorimotor noise (15,16). Common simplifications are neglecting feedforward control (21–24), using only local feedback control (23–28), and assuming that feedback gains are constant during large portions of the gait cycle (e.g. stance or swing) (21–28). However, some studies modelled the control policy by a deep neural network, which provides nonlinear full-state feedback (29–33). Few studies have explicitly modelled motor (22,24,26,27,34) or sensory (22) noise. Instead, most studies solve for control parameters that stabilise walking against numerical noise (i.e. accumulation of small integration and discretisation errors) or external perturbations (e.g. obstacles (21,32)). Usually, the parameters of the control policies are optimised via evolutionary strategies (21,22,24,28,34) or reinforcement learning (25,29–33). Some studies have leveraged the efficiency of trajectory optimisation methods (i.e. using direct collocation) to optimise control policies in the presence of noise. They either solve for a control policy that minimises a cost over a set of random samples from the noise distribution (26,35), or they represent the noise and state distributions by their mean and covariance (i.e. assume Gaussian distributions) (8,36). Few studies evaluated the effect of noise on the kinematics and/or control policy. Koelewijn et al. found that average foot clearance increased with the magnitude of the motor noise in a torque-driven biped (i.e. no muscle mechanics) with local proportional-derivative control (26). They also found that, in a musculoskeletal model with motor noise, co-contraction of thigh muscles is a minimal effort strategy for transtibial amputees walking with a prosthesis (27).

Both nonlinear musculoskeletal dynamics and sufficiently flexible control strategies are needed to study how sensorimotor noise influences walking, which poses numerical challenges. Methods that account for the effect of sensorimotor noise by random samples (taken either before (26,35) or during optimisation (22,24,26,27,34)) require large numbers of samples to be accurate. While conceptually simple, this approach would lead to high computational costs for models with many states and noise sources. Approximating the state distribution by a Gaussian distribution, which can be described by its mean and covariance, is computationally more efficient. The dynamics of the state covariance can then be described by Lyapunov differential equations based on a local first-order approximation of the system dynamics around the mean state (8,37). When the dynamics are highly nonlinear, the propagation of the state covariance will be inaccurate. The linearised dynamics could even display unphysiological behaviour, such as negative muscle activations or vertical ground reaction forces. The unscented transform (38,39) has been proposed as an alternative for local linearisation that can better handle highly nonlinear dynamics. This approach has been applied in optimal control simulations of aerospace applications (40,41) but not yet for simulations of biological movement.

Here, we present simulations of walking based on a neuromusculoskeletal model with 9 degrees of freedom actuated by 18 rigid-tendon Hill-type muscles that account for sensorimotor noise. The control strategy consists of time-varying feedforward and full-state feedback. We used a new approach, which combines a Gaussian approximation of the state distribution with an unscented transform to propagate the state covariance, direct collocation, and implicit dynamics to formulate the stochastic optimal control problem as an approximate deterministic problem. We evaluated how different levels of sensory and motor noise influenced the gait pattern. We compared the mean and variability of joint, centre of mass, and foot kinematics and muscle activations across simulated sensorimotor noise levels and with experimental data. We further examined whether variability in the simulated walking motion reproduces key features of experimentally observed kinematic variability. Experimental studies suggest that humans regulate task-level variables, such as centre-of-mass (COM) kinematics, more tightly than individual joint angles during walking (42–46). We therefore evaluated whether joint angle variability lies predominantly along the uncontrolled manifold of the COM position, i.e. co-variation of joint angles that does not influence COM position.

## Methods

### Neuromusculoskeletal model

We simulated walking based on a neuromusculoskeletal model (Fig 1) by assuming optimal control. We adapted a commonly used 2-dimensional musculoskeletal model (OpenSim’s gait10dof18mus (47,48)) by locking its lumbar joint in the anatomical reference position. The resulting model has 9 degrees of freedom: 3 at the floating base (pelvis forward, pelvis vertical, and Head-Arms-Torso tilt) and 1 at each hip, knee, and ankle. We used Newtonian rigid body dynamics to model skeletal motion (49,50). Each leg is actuated by 9 Hill-type muscle-tendon units (51,52). Muscle lengths and moment arms in function of joint angles were expressed as polynomials (49,53). To reduce the computational complexity, we assumed rigid tendons and ideal excitation-activation coupling (i.e. no dynamics). The simplified muscle mechanics do include length and velocity dependence of the contractile force and the effect of pennate muscle fibres. Each joint has viscous friction (coefficient 0.1 Nm s rad^-1^) and nonlinear stiffness to represent the net effects of ligaments (17,54). We modelled foot-ground contact as deformable spheres (foot) interacting with a rigid plane (ground). The force acting on each sphere was calculated from Hertzian stiffness and Hunt-Crossley dissipation (55,56). Each foot has two contact spheres: one at the heel and one at the forefoot.

**Fig 1.**
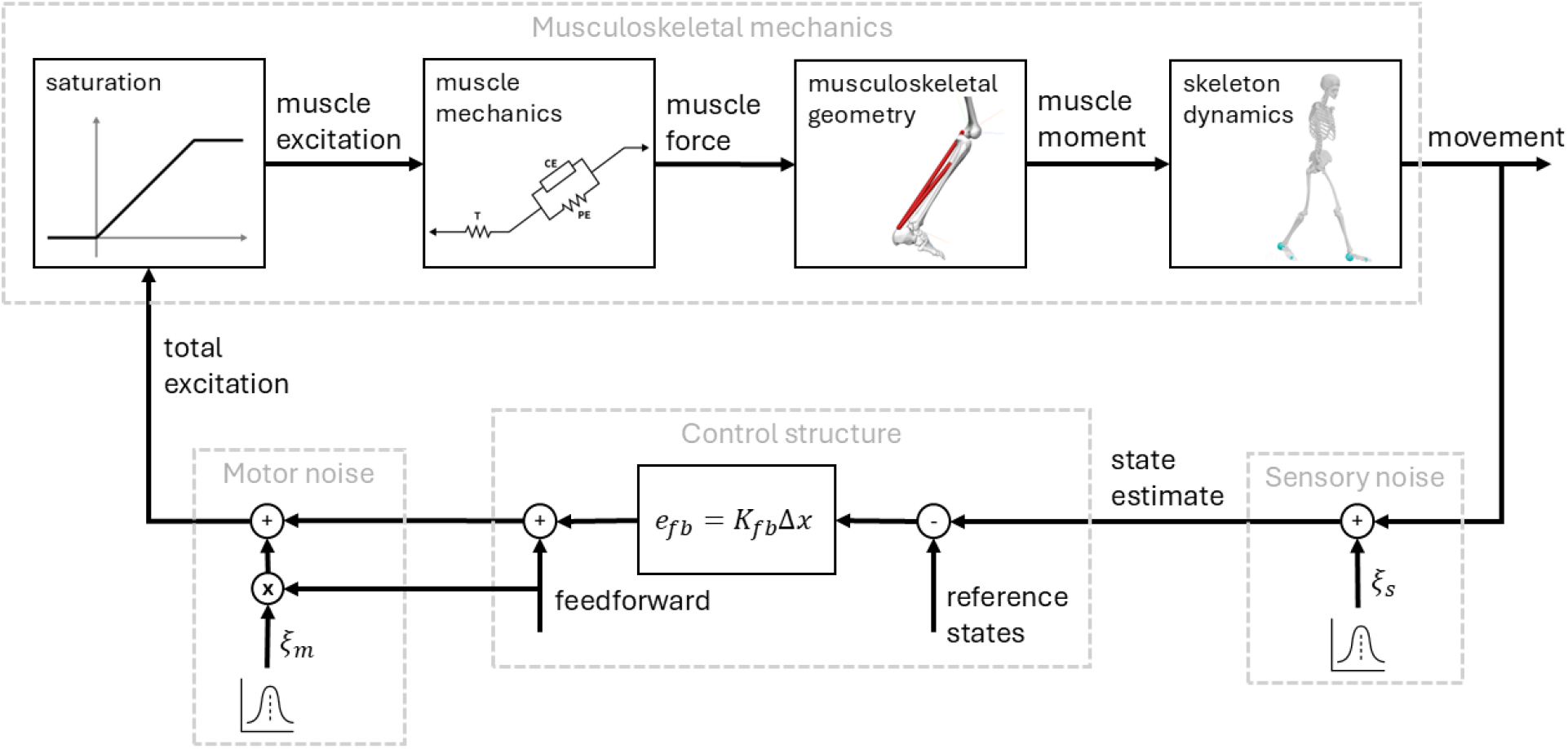
Block diagram of the neuromusculoskeletal model. Movement of the musculoskeletal system is controlled via feedforward and feedback muscle excitations. Feedforward excitations and feedback gains (***K***_***fb***_) are time-varying. The mean states are used as a reference for feedback. Muscle excitations are corrupted by signal-dependent, zero-mean Gaussian noise (i.e. the standard deviation of the noise scales with the mean total excitation, which is equal to the feedforward excitation). State estimates are corrupted by additive zero-mean Gaussian noise. Musculoskeletal illustrations were created with OpenSim (48).

We modelled neural control of muscles as the sum of feedforward excitations and linear full-state feedback (Fig 1). Feedforward excitations and feedback gains are time-varying. Feedback is based on the difference between reference states (mean states) and an estimate of the current state. We expressed the state estimate as the actual state corrupted by additive zero-mean Gaussian noise. This noise represents the lumped uncertainty due to sensory noise, delays, and internal model errors (57). The relative magnitudes of the uncertainty on state estimates (**Table 1**) are based on estimates of proprioceptive, visual, and vestibular noise (8). When redundant sensory information was available (i.e. pelvis and HAT), we assumed they were combined to form a maximum likelihood estimation (57).The absolute magnitude is determined by the sensory noise level (Ξ_s_). The control excitations are corrupted by signal-dependent Gaussian noise with a standard deviation that increases proportionally to the mean control (i.e. feedforward excitation) (1,2). The variance of this multiplicative noise is equal to the motor noise level (Ξ_m_). To ensure the muscle excitations remain above baseline (2%) and below maximum (100%), we applied a saturation function on the total excitations (See S1 Text for details). For example, if excitation is low, inhibitory feedback is still possible but will not reduce muscle excitation below the baseline level. As a consequence of introducing noise, the state is stochastic and should be described by a probability distribution.

**Table 1.**
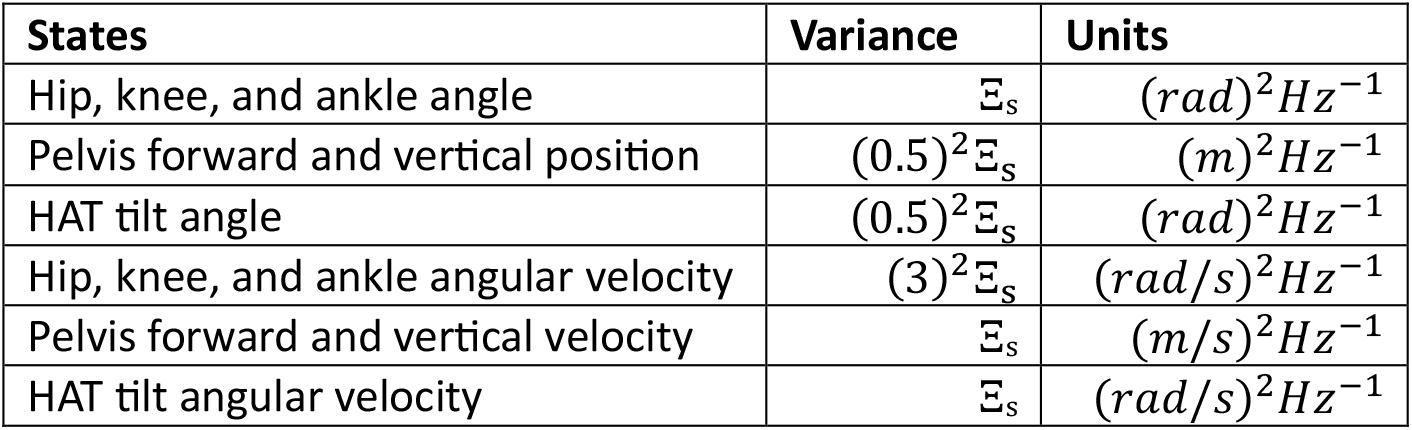
Magnitude of the variance of state estimates. Relative magnitudes are based on estimates of proprioceptive, visual, and vestibular noise (8), geometry, and an optimal combination of redundant information. The level of sensory noise (Ξs) varied between different simulations (range: 0 – 0.02^2^).

| States | Variance | Units |
| --- | --- | --- |
| Hip, knee, and ankle angle | $\Xi_s$ | $(rad)^2 Hz^{-1}$ |
| Pelvis forward and vertical position | $(0.5)^2 \Xi_s$ | $(m)^2 Hz^{-1}$ |
| HAT tilt angle | $(0.5)^2 \Xi_s$ | $(rad)^2 Hz^{-1}$ |
| Hip, knee, and ankle angular velocity | $(3)^2 \Xi_s$ | $(rad/s)^2 Hz^{-1}$ |
| Pelvis forward and vertical velocity | $\Xi_s$ | $(m/s)^2 Hz^{-1}$ |
| HAT tilt angular velocity | $\Xi_s$ | $(rad/s)^2 Hz^{-1}$ |

To find control laws (i.e. feedforward excitations and feedback gains) that generated walking, we assumed optimal control while imposing task constraints. We minimised the expected value of squared muscle activations (*a*), excitations (*e*_*tot*_), and joint limit torques (τ_*lim*_).

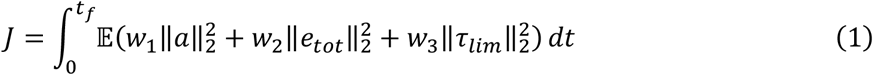

Squared muscle activations and excitations represent effort. Including squared joint limit torques penalises excessive loading of ligaments and joint capsule, which can be interpreted as avoiding pain. Previous simulation studies found that penalising muscle effort (here modelled as the sum of squared activations) is important to predict human-like gait (15,16). Therefore, we used weights *w*_*1*_ *= 1000, w*_*2*_ *= 0*.*02*, and *w*_*3*_ *= 0*.*1 (Nm)*^*-2*^. Preliminary simulations indicated that the values of *w*_*2*_ and *w*_*3*_ have a limited effect on the result, as long as they are much smaller than *w*_*1*_. To simulate locomotion, we imposed that the mean forward pelvis position equals *0* at the initial time and 0.6667 m at the final time (0.5556 s). These correspond to a single step with forward velocity 1.2 m/s and stride frequency 0.9 Hz, which is representative of an average healthy adult. All other mean states were constrained to be periodic and left-right symmetric without imposing specific values. For example, the initial left ankle angle should equal the final right ankle angle, the initial and final vertical pelvis velocity should be equal, etc. We used the same mapping to constrain the initial and final state covariance matrix (including forward pelvis position). Since we imposed periodicity on the state distribution, the uncertainty cannot grow over time. Thus, only a stable limit cycle satisfies the task constraints.

### Stochastic optimal control

The resulting optimal control problem is stochastic due to the presence of noise. It has highly nonlinear dynamics with 18 states and 36 noise sources, making it a very challenging problem. To solve this stochastic optimal control problem, we approximated the state distribution by a Gaussian distribution. A Gaussian distribution is fully determined by its mean and covariance. To describe how the state mean and covariance change over time, we combined the unscented transform with implicit formulation of dynamics and direct collocation. The unscented transform was developed as a method to propagate the mean and covariance through a nonlinear transformation (e.g. nonlinear dynamics) without requiring a linear approximation of the transformation (38,39). It is based on the notion that the mean and covariance of a distribution can be represented exactly by a finite set of (well-chosen) samples (referred to as sigma points). The transformed distribution can be estimated by first applying the nonlinear transformation to each sample and then calculating the weighted mean and covariance of the transformed samples (38,39). Ross et al. proposed an approach where the nonlinear transformation maps the initial states to the states at the end of the time horizon (58). The sigma points are sampled from the distribution of the initial states (58) or the distribution of uncertain parameters of the dynamics (41). This results in a set of state trajectories with a single control strategy, which can be found via trajectory optimisation methods (41). To include process noise (such as sensorimotor noise), the noise injected during each discrete time step should be included in the distribution from which the sigma points are sampled. This would result in thousands of sigma points and state trajectories. Ozaki et al. proposed a method that handles process noise more efficiently (40). They considered a nonlinear transformation that maps the states from one mesh point to the next, i.e. one time step with an explicit integrator. Sigma points are then sampled from the distribution of states and noise injected during one discrete time step. At each mesh point, the transformed sigma points are combined into a mean and covariance from which sigma points for the next time step are sampled (40). Hence, the states at a mesh point are expressed as a function of all previous controls and the initial state. Using this to formulate an optimal control problem, corresponds to using a direct single shooting approach. In musculoskeletal simulations, it has been demonstrated that using direct collocation instead of direct shooting (59,60) and implicit formulation of dynamics (51,61) greatly improves the efficiency of the simulations and facilitates using complex models. Hence, applying the approach presented by Ozaki et al. to musculoskeletal models is not expected to work well. Therefore, we developed a new approach which combines the unscented transform with direct collocation and implicit formulation of dynamics.

The presented approach is not tied to the chosen neuromusculoskeletal model or control law. Therefore, we first describe it in the general case. Symbols and notations are summarised in Table 2, matrices are in bold. The time horizon is discretised in *N*_*m*_ mesh intervals (here, *N*_*m*_ *= 64)*. On each mesh point, the state distribution is approximated by a Gaussian distribution. We introduced optimisation variables to represent the mean state 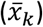 and the Cholesky factor of the state covariance matrix (***L***_***k***_ ; i.e. the lower triangular matrix ***L***_***k***_ such that the state covariance matrix 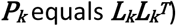 at every mesh point and the controls over every mesh interval (*u*_*k*_). In each mesh interval, we used the unscented transform to propagate the mean state and state covariance over the mesh interval. First, the initial state distribution is augmented to include noise sources (38), augmented state variables are indicated by left superscript a:

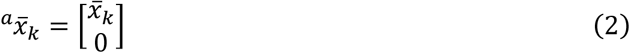

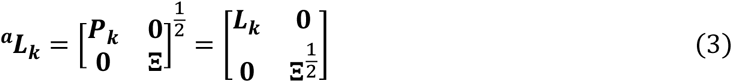

where **Ξ** is the noise covariance matrix, which is determined based on the level of sensory and motor noise. From this augmented distribution, *N*_*σ*_ sigma points (light blue diamonds in Fig 2) are sampled. The first sigma point lies at the mean, the others on the c^th^ covariance contour (here, *c = 3*). The (augmented) state vectors at all sigma points, combined in the matrix ***χ***_*k*_, are then computed as

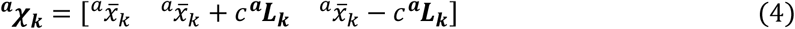

**Table 2.**
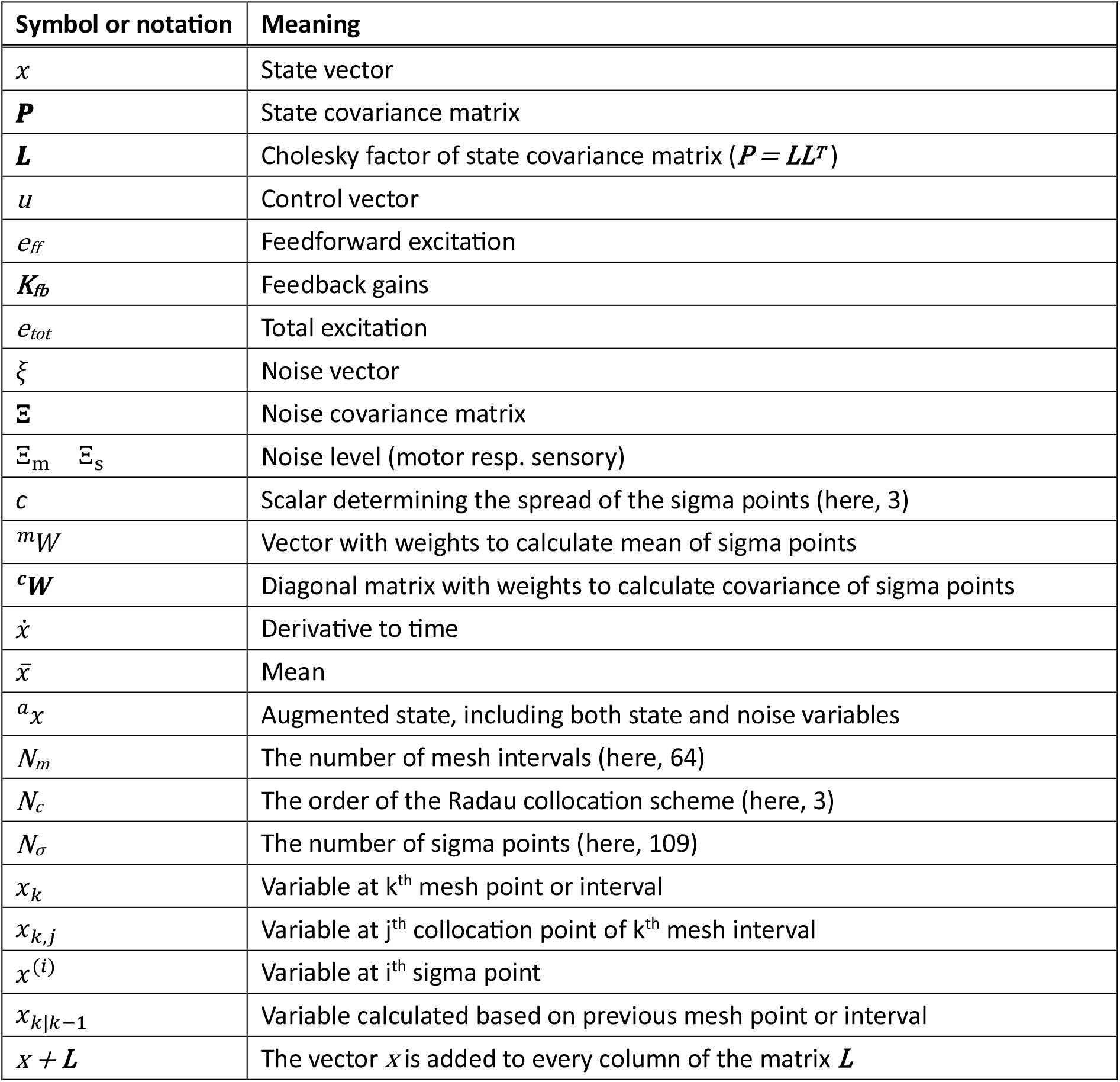
Summary of symbols and notations.

**Fig 2.**
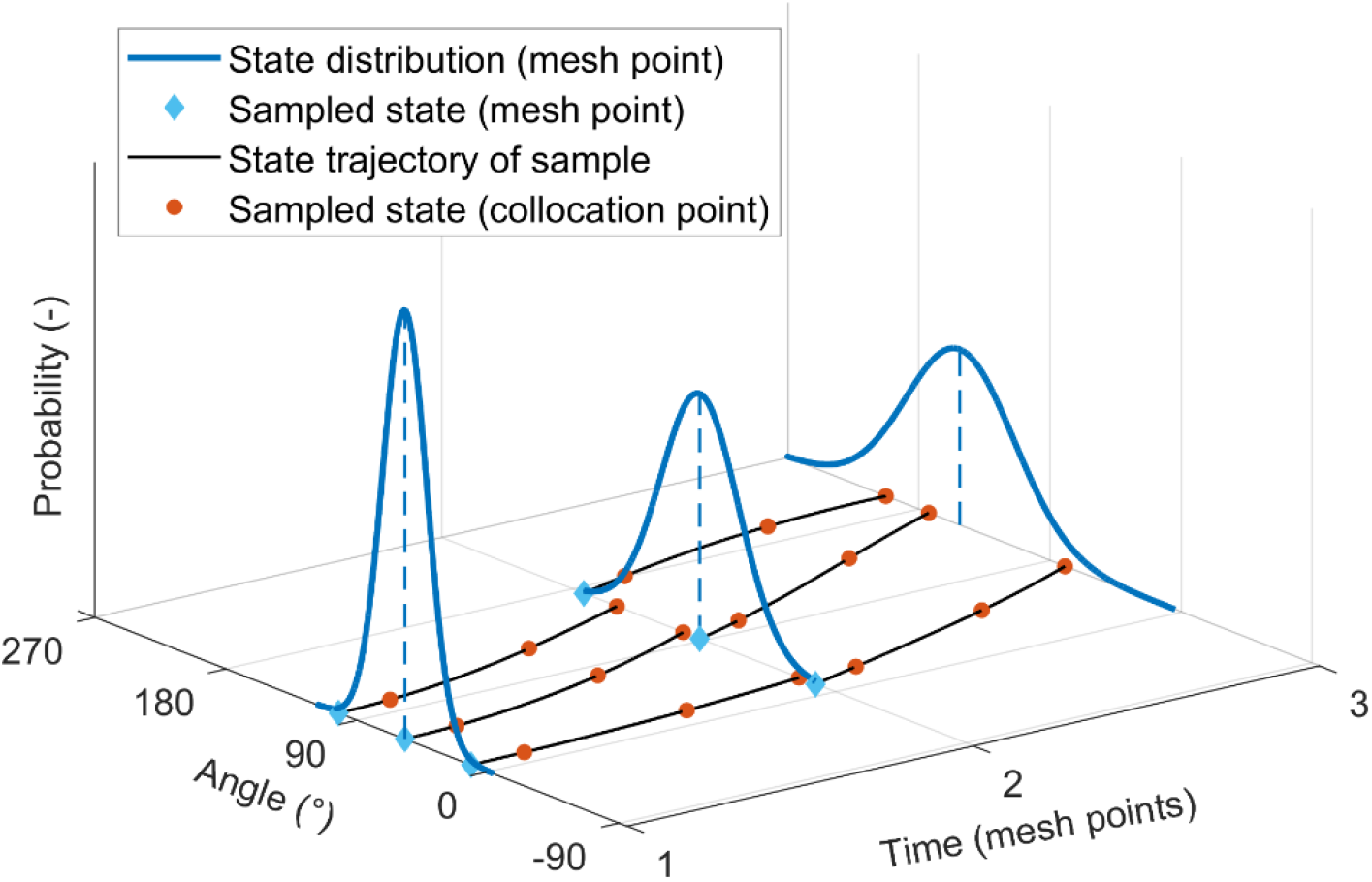
Illustration of the proposed stochastic optimal control approach. At every mesh point, sigma points are sampled from the state distribution. For each sigma point, the state trajectory over the mesh interval is described using direct collocation (and implicit dynamics). At the end of the mesh interval, the state mean and covariance are computed based on the sigma point states. The mean and covariance are imposed to equal the state distribution at the next mesh point.

Since the noise distribution is not time-varying, the sampled noise vectors (ξ) are the same in every mesh interval:

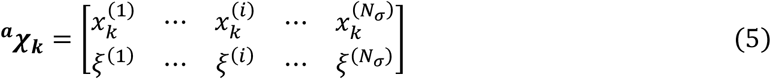

where the right superscript is the index of the sigma point (i). For each sigma point, the state trajectory (black lines in Fig 2) is represented by *N*_*c*_ Radau collocation points (here, *N*_*c*_ *= 3*; orange dots in Fig 2). The state vector and its time derivative 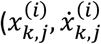 subscript j is the index of the collocation point) at every sigma-collocation point are optimisation variables. The collocation equation

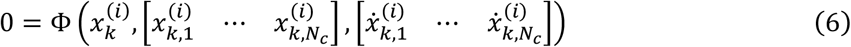

is added as path constraint. At every collocation point, the implicit system dynamics

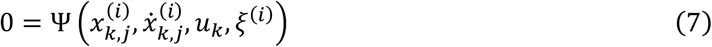

are added as path constraints. The state vectors at the next mesh point, based on the current mesh interval (subscript k+1|k), are

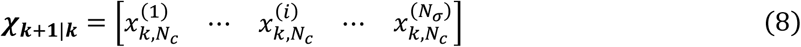

because in a Radau collocation scheme the last collocation point coincides with the next mesh point. The mean state and state covariance at the next mesh point can be calculated as the weighted mean and covariance of the states at the sigma points. The derivation of the weights (vector ^m^W and diagonal matrix ^**c**^**W**), based on the value of *c* in equation 4, is available in S1 Text.

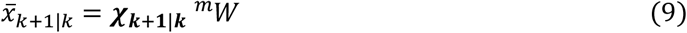

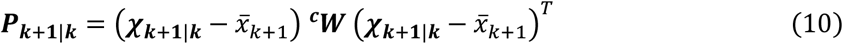

The state distribution at the next mesh point is also represented by the optimisation variables 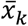 and ***L***_***k***_ . To ensure continuity of the state distribution, both expressions should describe the same mean State

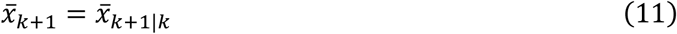

and the same and state covariance

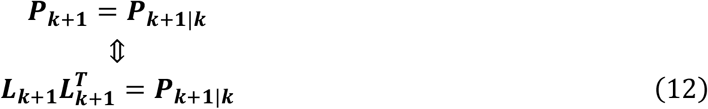

Substituting equation 9 into equation 11 and equation 10 into equation 12, yields

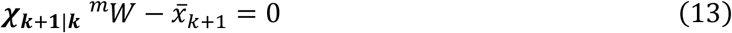

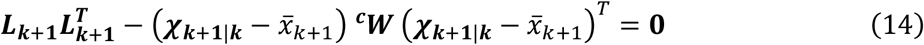

which are added as path constraints to impose continuity on the state distribution. The expected cost over a mesh interval can be calculated as the weighted mean of the cost associated with each sigma point. Consider a cost y_k_, which can represent any of the cost terms from equation 1 evaluated over the k^th^ mesh interval for a sigma point.

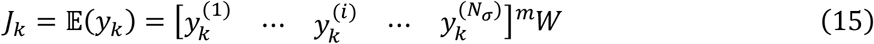

To impose periodicity and left-right symmetry, constraints linking the mean state and state covariance at initial and final time are added

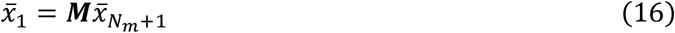

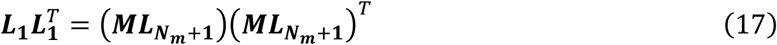

with ***M*** a permutation matrix that encodes left-right symmetry of the states. When imposing periodicity without symmetry, ***M*** is the unit matrix.

We introduced the total excitation (*e*_*tot*_) as optimisation variables at each sigma-collocation point, as preliminary simulations and analysis indicated that this improved the numerical properties of the problem. The control law is then imposed by added the following path constraint

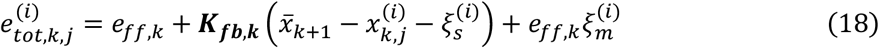

As a result, the expected excitation effort (equation 1) can be expressed in two different ways. First, the expected value of the squared excitation can be calculated via equation 15 and the left-hand side of equation 18. Second, it can be calculated based on the right-hand side of equation 18, which linearly depends on states and noise.

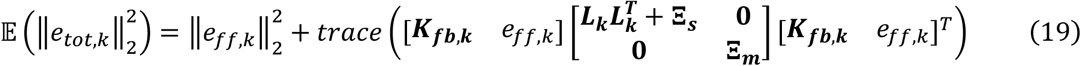

In the solution, both expressions for the expected excitation effort are equivalent. We included both expressions in the objective function The path constraints depend only linearly on e_ff_, so the variable needs to be included in the objective function to avoid a singular arc. We also observed that having e_tot_ and K_fb_ in the objective function improved the convergence of the solver. In summary, including both expressions in the objective function improved the numerical properties of the problem without needing additional regularisation terms.

The simulations were implemented with CasADi (62), using its interface to MATLAB (The Mathworks Inc., USA). We used 64 mesh intervals and a third order Radau collocation scheme. The nonlinear program was solved with ipopt (63) and ma97 (64,65). Derivatives were obtained using algorithmic differentiation with CasADi (62). These derivatives are exact, as opposed to finite difference approximation, which improves the convergence of the simulation (49). We scaled the constraints such that the acceptable violation of all constraints had a similar magnitude. We scaled the optimisation variables such that the nonzero elements in the constraint Jacobian (i.e. the sensitivities of the constraints to the optimisation variables) had similar magnitude, as this improves the numerical stability of the solver algorithm (63,66). All simulations were performed on a desktop computer with an *AMD Ryzen Threadripper Pro 7975WX* CPU (32 cores, 4 GHz) and 512 GB DDR5 RAM, running Windows 11.

To reduce the computational cost, most calculations were offloaded to compiled code instead of being evaluated in CasADi’s built-in virtual machine (62). The model had 18 states and 36 noise sources, thus requiring 109 sigma points. At each of the resulting 20928 collocation points, the neuromusculoskeletal dynamics and their derivatives need to be evaluated. We implemented the dynamics in MATLAB (cf. previous work (53)), then used CasADi’s built-in code generator (62) to create C code with functions to evaluate the path constraints (equations 6, 7, and 18) and their Jacobian. The code was then compiled and imported back into the nonlinear program. This reduced the evaluation time by a factor 20 for the dynamics constraints and by a factor 450 for their Jacobian. The path constraints on state covariance (equation 14) consist of multiplications and additions with dense matrices. The computational cost of evaluating these scales with the size of the matrices (i.e. the number of states) to the third power, thus, could pose a bottleneck for using more complex models. To speed up the evaluation, we relied on a Basic Linear Algebra Subprograms (67) library that was optimised for our hardware (68). This reduced the evaluation time by a factor 9 for the covariance constraints and by a factor 4 for their Jacobian. We used parallel computing to further reduce simulation time. Evaluation of constraints and their Jacobian was spread over 8 CPU threads (8 mesh intervals per thread), ma97 was assigned to use 16 CPU threads. More details and benchmarks are given in S2 Text.

### Analysis of simulation results

We performed a series of simulations to evaluate the effect of noise on the gait pattern. From eight sensory noise levels (Ξ_s_: 0^2^, 0.001^2^, 0.002^2^, 0.003^2^, 0.004^2^, 0.005^2^, 0.010^2^, 0.020^2^) and three motor noise levels (Ξ_m_: 0.005^2^, 0.010^2^, 0.020^2^), we selected 16 combinations. We also performed a simulation without noise (deterministic). We did not simulate the effect of sensory noise in the absence of motor noise because the noise can be negated entirely by not using feedback, thus it is equivalent to the deterministic case.

We evaluated predicted muscle activities and movement patterns (mean and variability) against experimental data from 18 young adults walking on a treadmill at 1.1 m/s (69). To prepare the experimental data for comparison with simulation results, we performed additional processing. First, we scaled the data of each subject. Filtered electromyography (EMG) signals were scaled relative to their peak value, ground reaction forces (GRF) were scaled to body weight, vertical pelvis position was scaled to leg length, and forward pelvis position was offset such that the mean over the trial was zero. Second, every gait cycle (starting at right side heel-strike) was interpolated to 129 time points (cfr. 64 mesh intervals per step). At each time point, we calculated the mean and standard deviation across the different gait cycles (on average, 62 per subject). For EMG and GRF, we also calculated the median and 99.7 % confidence interval (cfr. simulated sigma points lie on the 3^rd^ covariance contour). Third, we calculated the group average of mean, standard deviation, median, and confidence interval of variables.

Because experimental studies indicate that the centre of mass (COM) position is tightly controlled during walking, we included two composite outcome measures to analyse sagittal plane centre of mass kinematics: the COM angle and the ratio of variance that lies along the uncontrolled manifold over the total variance. Inspired by the inverted pendulum model of gait, we computed the COM angle as the angle of the line between the heel and the whole-body centre of mass, relative to the vertical direction. We used the uncontrolled manifold method as described in (42,43) to evaluate whether joint position noise in experiments and simulations was similarly structured. We expressed the COM position as a function of foot-to-ground, ankle, knee, and hip angle (kinematic chain), derived the Jacobian, and computed the null-space of this Jacobian, i.e. the uncontrolled manifold. Then, we decomposed the variability of the four angles into a variance parallel to 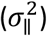 and normal to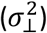the uncontrolled manifold of the COM position. The former represents the variability in the angles that does not affect the COM variance, and the latter represents the variability that does. To compare the two components, we calculated the ratio of the parallel to total variance and scaled and centred it to obtain a variable between -1 and 1 (42):

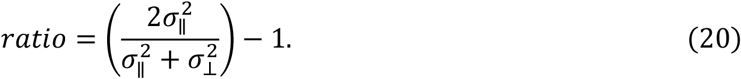

Hence, a value of zero means equal variance along and normal to the uncontrolled manifold, whereas a positive value means larger variance along the uncontrolled manifold (i.e. not influencing COM position).

## Results

We performed stochastic optimal control simulations of walking based on a 2D model with 9 degrees of freedom and 18 muscles to study how the level of sensory and motor noise influences the control strategy and gait pattern, given the same neuromusculoskeletal dynamics and performance criteria (i.e. minimising expected effort). From three levels of motor noise and eight levels of sensory noise, we selected 16 combinations. Finding an optimal control policy for a sensorimotor noise level took, on average, 65 hours (range 14 – 254 hours; elapsed real time). We also performed a deterministic simulation for the case without any noise. Here, optimising the muscle excitation patterns took 14 seconds. We compared kinematics, muscle activations, ground reaction forces, swing foot clearance, and centre of mass kinematics between simulations with different noise levels and experimental data. Time traces of all simulation results are available in S3 Text. Each combination of sensory and motor noise has its own colour (see Fig A in S3 Text for colour scheme), which is used across figures.

Accounting for sensorimotor noise had little effect on the mean kinematics, i.e. mean kinematics of the stochastic simulations were very close to the mean kinematics of the deterministic simulation (Fig 3a, Fig B-H in S3 Text). The mean simulated kinematics were similar across most sensorimotor noise levels and captured key features of human walking, i.e. pelvis moves up and down twice per stride, the hip goes to extension then flexion, knee is extended during stance and flexed during swing, and ankle kinematics shows the three rockers (plantarflexion during initial contact, dorsiflexion during mid-stance, plantarflexion during terminal stance). Notwithstanding this general agreement, there are considerable differences between simulated and experimental kinematics, especially for the hip, knee, and ankle, between 30 and 50% of the gait cycle. These differences can be attributed to the use of a relatively simple model with rigid tendons as our simulations based on a 3D model with compliant tendons do capture sagittal plane kinematics throughout the gait cycle (18). At high sensory noise levels (0.01^2^ and 0.02^2^), mean kinematics showed increased hip and knee flexion and ankle plantarflexion during the swing phase (Fig B-H in S3 Text).

**Fig 3.**
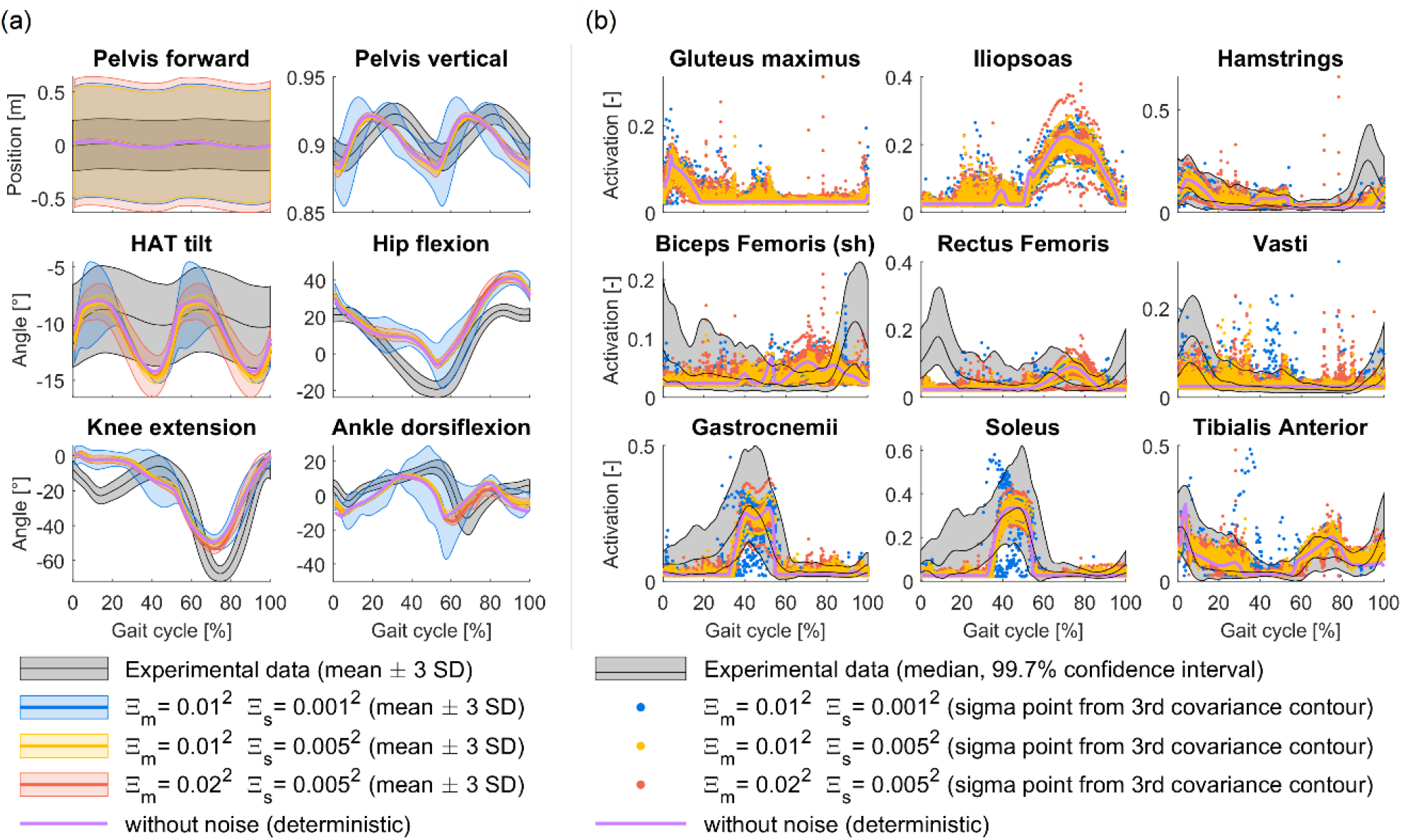
Simulated and experimental kinematics and muscle activations. All graphs represent the 99.7% confidence interval. Joints and muscles from the left side are omitted due to symmetry. (a) Kinematics were assumed to follow a Gaussian distribution, hence are given by the mean and standard deviations (SD). (b) Muscle activations are evaluated in the sigma points at the end of each mesh interval (Fig 2).

Sensorimotor noise levels strongly affect the kinematic variance. We used the trace of the state covariance matrix averaged over time as a summary measure for kinematic variance (Fig 4a). Variance increased with motor noise level and changed non-monotonically with sensory noise level. Variance was highest at low or high sensory noise levels and was lower at intermediate sensory noise levels. We obtained the lowest variance in the absence of sensory noise. We found the same relationship between averaged variance and sensorimotor noise levels when only considering angular positions or angular velocities (Fig I in S3 Text).

**Fig 4.**
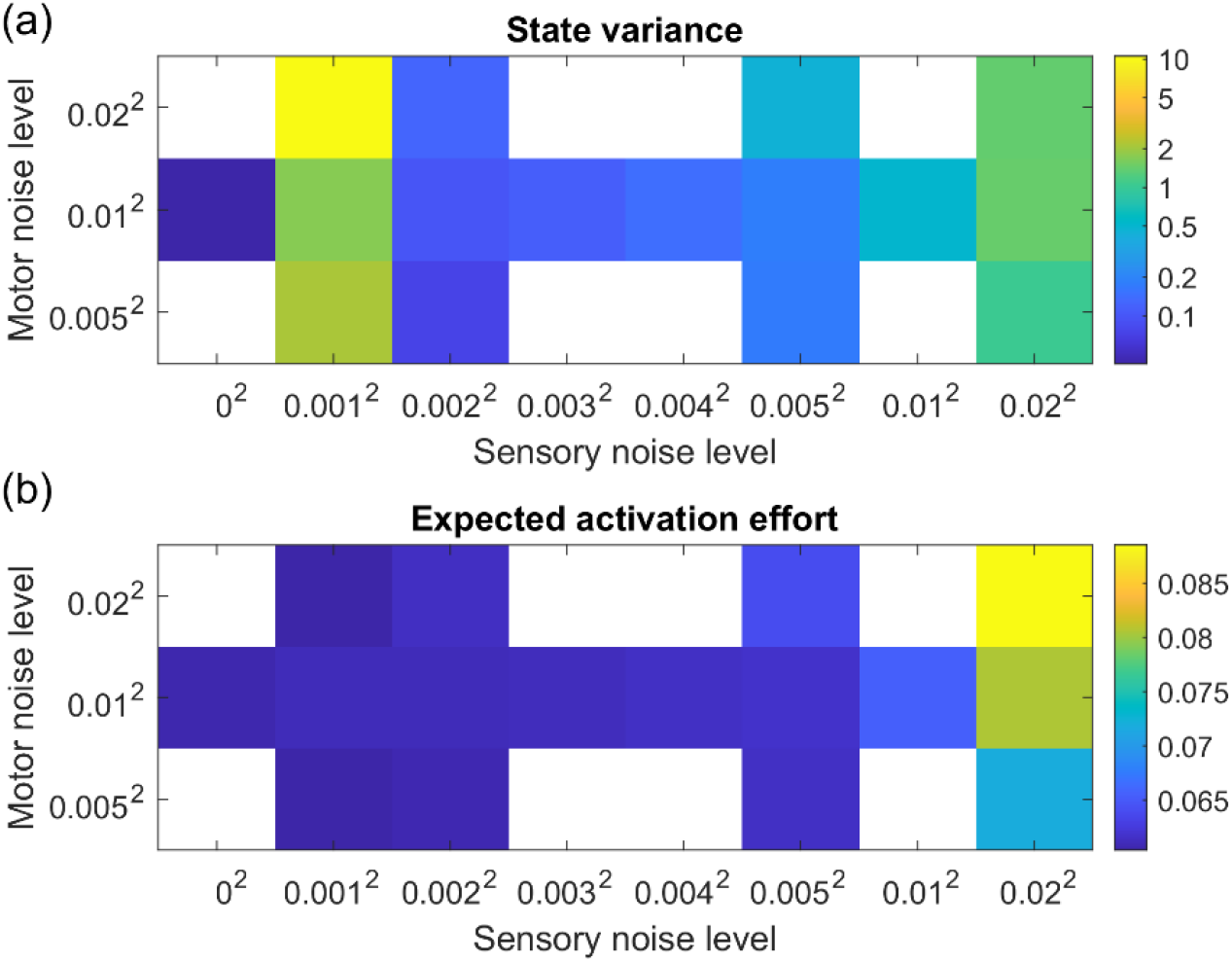
State variance and expected effort for different noise levels. Each square represents the result of a simulation with given sensory and motor noise levels. We did not simulate all combinations of noise levels. (a) State variance was calculated as the trace of the state covariance matrix at each mesh point, averaged over the time horizon. (b) Expected activation effort is the major term in the objective that is minimised. It is calculated as the expected value of the sum of squared muscle activations, integrated over the time horizon.

We found that the magnitude of the simulated and experimental position standard deviations (SD) agreed best for simulations with low sensory noise (0.001^2^; Fig 5a, azure). However, we found that the normalised cross-correlation (NCC) between simulated and experimental position SD (shape) agreed best for the simulation with intermediate sensory and motor noise levels (NCC: 0.94 – 0.98; Fig 5a, amber). Yet, these simulations underestimated the magnitude of the position SD.

**Fig 5.**
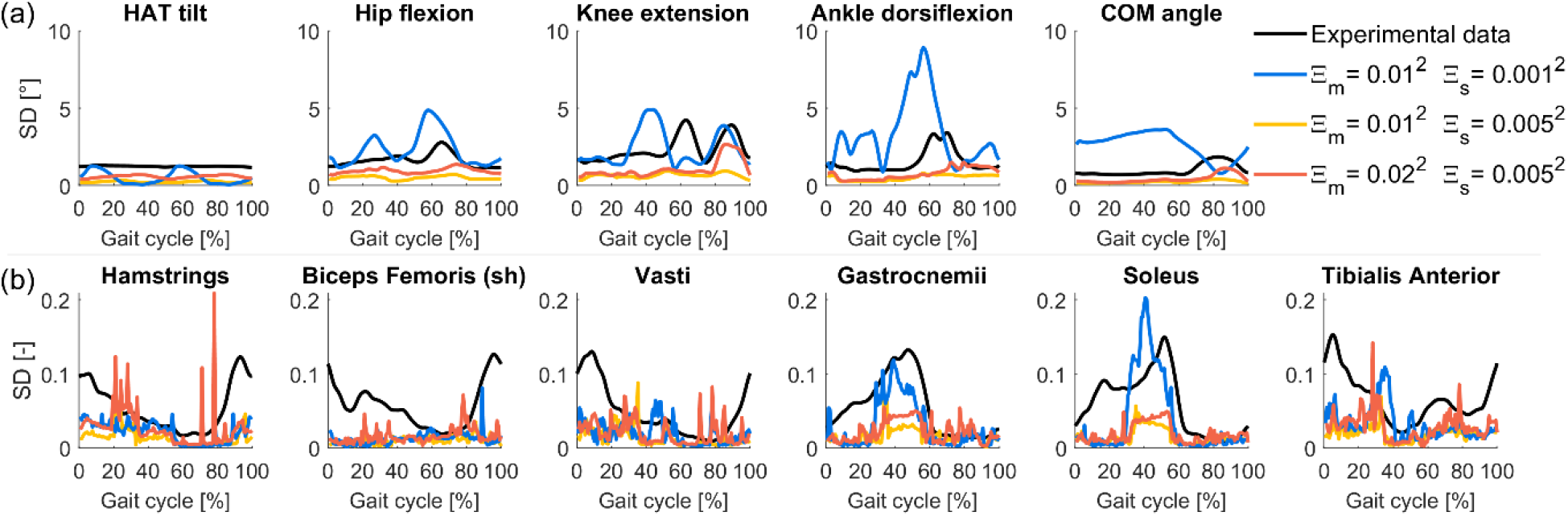
Comparison of standard deviations. (a) Selected (joint) angles. (b) Selected muscle activations

Simulated peaks in muscle activity are in agreement with experimentally observed peaks in muscle activity, whereas the agreement between experimental and simulated muscle activation variability depends on the sensorimotor noise level. Our simulation approach does not make assumptions about the distribution of the variables that aren’t states, such as muscle activations. Therefore, we evaluated muscle activations in each sigma point at the end of each mesh interval (Fig 2) and visualised these values. In other words, we did not attempt to estimate a mean and covariance based on these values, as their distribution is not necessarily Gaussian. Whereas the general agreement between simulated and experimental muscle activity patterns is good, the simulations fail to capture some experimentally observed peaks in muscle activation (Fig 3b). In particular, the simulations do not capture experimental hamstrings and biceps femoris short head activations during late swing (85-100% of the gait cycle) and experimental rectus femoris and vasti activation during early stance (0-15% of the gait cycle). The absence of simulated vasti activation is due to the absence of knee flexion during stance in this simple model (70) whereas it is likely that rectus femoris EMG is corrupted by cross-talk from vasti as simulated rectus femoris activity patterns resemble experimental observations from wire EMG (71). In addition, the onset of simulated gastrocnemius and soleus activity is delayed with respect to experimental observations, likely due to the assumption of a rigid tendon (18).

The estimated magnitude (based on the assumption of a Gaussian distribution) of muscle activation SD agreed best with experimental data for the simulation with low sensory noise and intermediate motor noise (Fig 5b, azure), similar to the agreement in position SD. In agreement with experimental observations, SD was higher when muscle activity was higher. However, simulations failed to capture the pattern of muscle activity SD and this could not be fully explained by differences in simulated mean muscle activity. In addition, simulations overestimated muscle activity SD of some muscle groups in mid stance (e.g. tibialis anterior; Fig 5b).

Expected effort monotonically increased with sensory noise level and in most cases also with motor noise level (except for the lowest sensory noise levels; Fig 4b). We used the sum of expected muscle activations squared as a measure of effort, hence both mean muscle activity and muscle activity variability shape effort but in a non-linear way due to the saturation (no activations below 0.02 or above 1). We found that the mean muscle activity slightly increased with sensorimotor noise levels with a steeper increase at the highest sensory noise levels (Fig 3b, Fig J in S3 Text). The variability of muscle activity increased with motor noise level and was highest at the lowest and highest sensory noise levels, similar to state variance.

Sensorimotor noise level had little effect on the mean ground reaction forces (GRF) or the duration of the stance phase (Fig 6a). Only at high sensory noise levels (0.01^2^ - 0.02^2^), the initial peak in GRF was higher. Compared to experimental observations, simulated GRF showed a steeper increase and higher initial peak after initial contact (0-15% GC). All simulations underestimated the duration of the stance phase. Similar to state variability, GRF variability was high at low and high sensory motor noise levels but low at intermediate sensorimotor noise levels.

**Fig 6.**
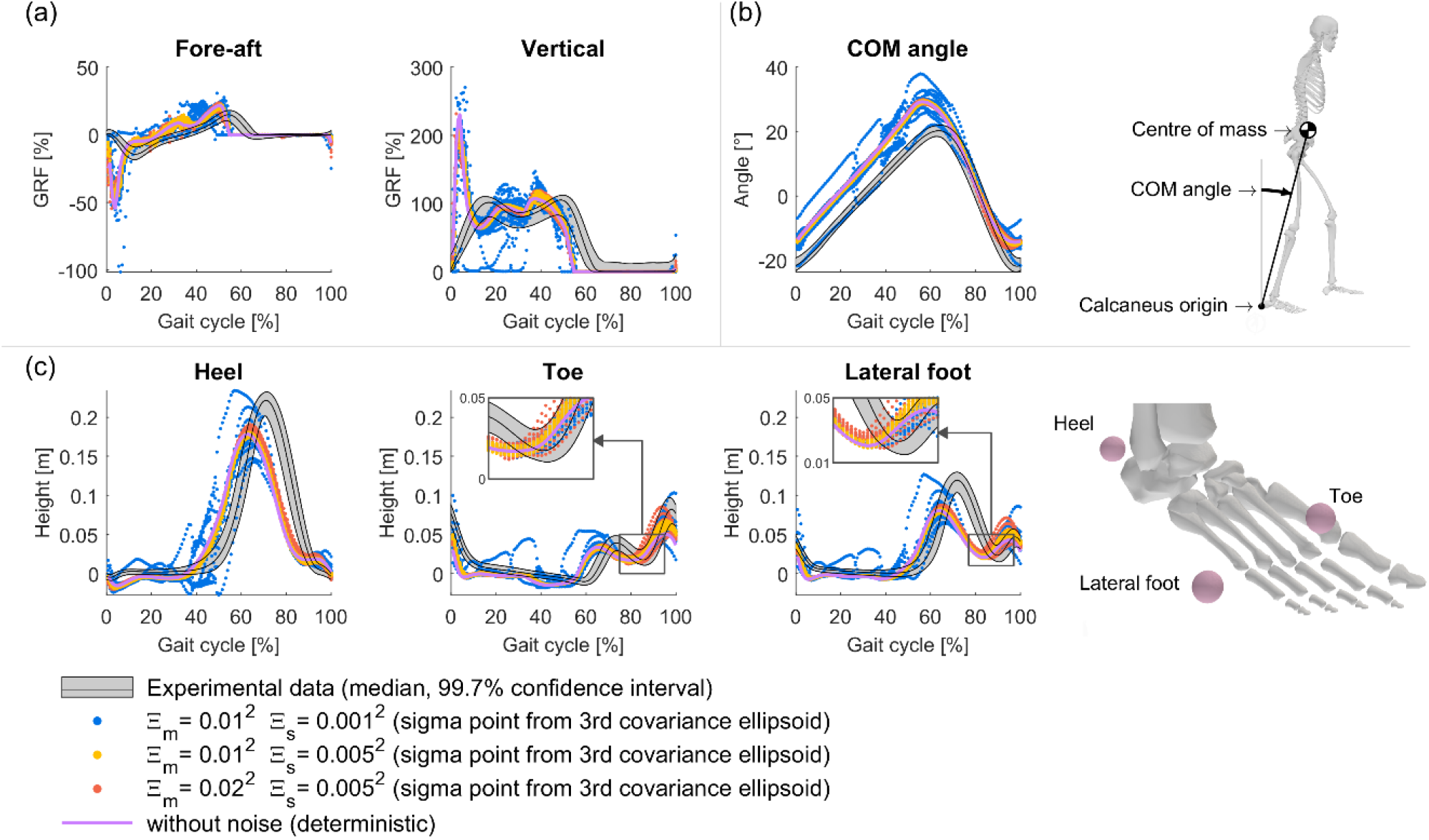
Simulated and experimental ground reaction forces, COM angles and foot marker heights. All simulated variables are evaluated in the sigma points at the end of each mesh interval (Fig 2). Musculoskeletal illustrations were created with OpenSim (48). (a) Ground reaction forces expressed as a percentage of body weight. (b) Angle of the approximating inverted pendulum. (c) Vertical position of markers on the foot, relative to their position during standing. Positive values represent foot clearance, negative values represent deformation of the foot and shoe.

Sensorimotor noise level had little influence on minimal foot clearance but affected mean foot clearance and variability of foot clearance (Fig 6c, Fig K in S3 Text). We characterised foot clearance by the vertical position of three foot markers, offset by their position during standing. Positive values represent foot clearance, negative values represent deformation of the foot and shoe. Simulations resulted in marker heights that captured the overall shape and magnitude of the experimental curve, except for underestimating peak foot clearance. Timing was shifted due to the shorter stance time in the simulation. The mean and variability change with noise level, where foot clearance variability was larger when the state variance was larger. Yet, differences in variability across sensorimotor noise level seemed smaller for foot clearance than for state variance (moment of minimal foot clearance at 80-90% GC, visual inspection). Across noise levels, during swing (60-95% GC), the foot markers were lifted by at least 1 – 2 cm at all sigma points (3rd covariance contour). In the deterministic simulation, the foot markers were lifted by at least 2 cm. These minimum heights of the toe and lateral foot marker in swing (80-90% GC) were in agreement with experimental data.

Sensorimotor noise level did not affect the mean centre of mass angle trajectory (Fig 6b). The COM angle was defined as the line between the calcaneus origin and the whole-body centre of mass, relative to vertical. The mean COM angle increased linearly while the foot was on the ground. Simulations captured the mean curve very well, given a constant offset of 7° (COM is further forward). The mean was not affected by noise level, while the variability increased with increasing state variance. Most simulations captured that COM angle variability was larger in swing than in stance, but underestimated the magnitude (NCC: 0.99; Fig 5a). At a low sensory noise level (0.001^2^), simulations overestimated the magnitude and did not capture how the variability of the COM angle changed over the gait cycle.

In agreement with experimental observations, the simulations predicted that joint angle variability around the step-to-step transition had a limited effect on COM position (Fig 7b). We evaluated the ratio of variance parallel to and normal to the uncontrolled manifold (i.e. joint angle subspace that does not affect COM position). For most sensorimotor noise levels and in agreement with experimental data, the simulated ratio is high at initial contact, drops to a low value (negative) during mid-stance (10 – 70 % stance), and is high at the end of stance (Fig 7a). However, for most subjects, the ratio reaches its minimum value earlier in stance than in simulations. In addition, simulations with low sensory noise do not capture the high ratio at heel contact.

**Fig 7.**
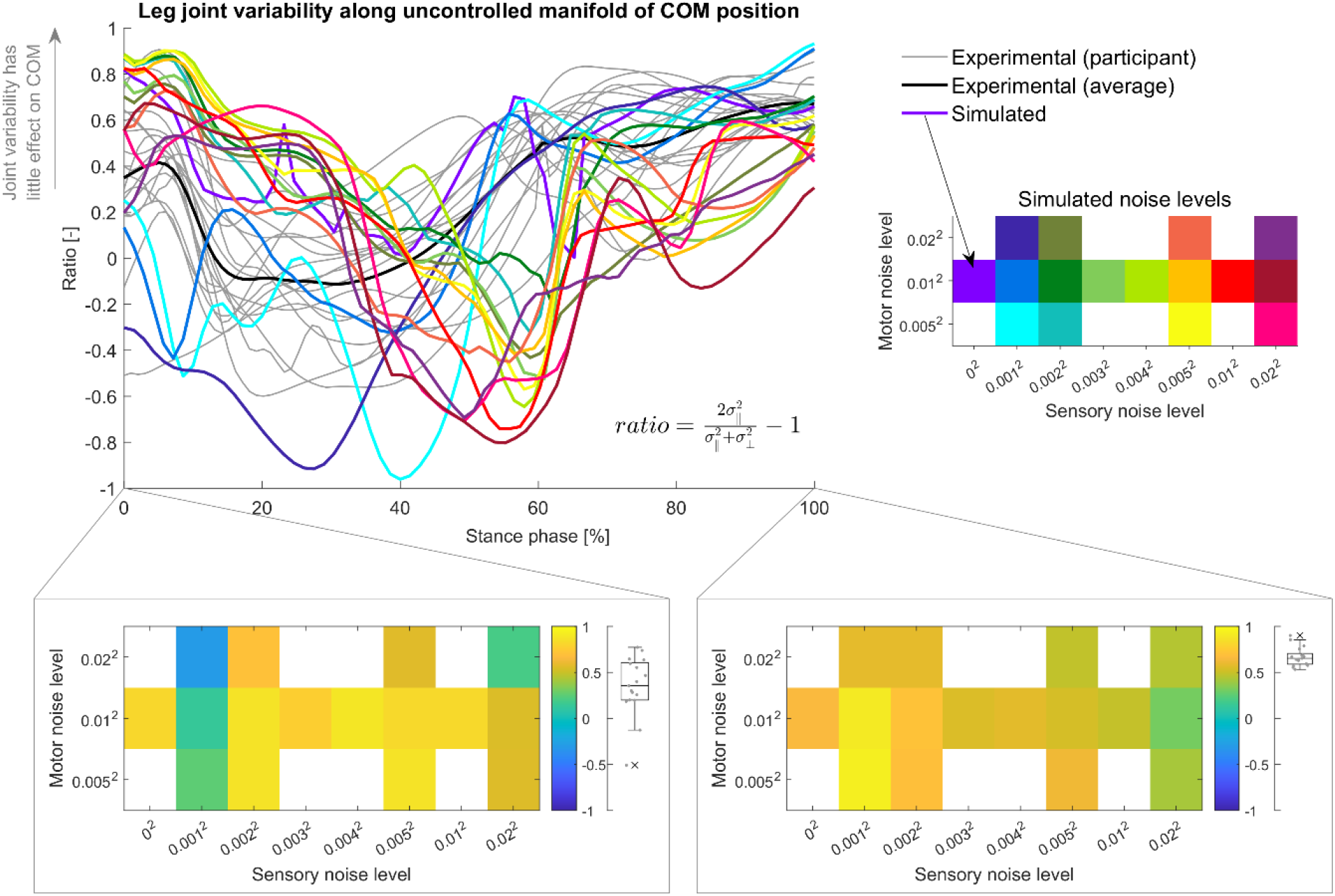
Uncontrolled manifold analysis of centre of mass position control. We expressed the horizontal and vertical centre of mass (COM) position as a function of foot-to-ground, ankle, knee, and hip angle (kinematic chain). Then, we decomposed the variability of the four angles into a variance parallel to and normal to the uncontrolled manifold of the COM position and computed the ratio between the variance parallel to the uncontrolled manifold and the total variance. The top panel shows the ratio during the stance phase. The bottom panels show the ratio at initial and final stance (Simulation results on grid, experimental data in boxplot).

## Discussion

We performed stochastic optimal control simulations of walking based on a neuromusculoskeletal model that was considerably more detailed than previously used models. Our numerical approach was based on approximating the state distribution by a Gaussian but in contrast to previously proposed approaches (8) used an unscented transform to propagate the distribution, which is more robust against nonlinearities than methods based on local linearisation. The ability to perform stochastic optimal control simulations based on detailed models, allowed us to study the effect of sensorimotor noise on walking without making strong assumptions on the structure of the controller. We modelled feedforward as well as feedback control, where feedback was based on uncertain information on the full state. The simulated mean kinematics and muscle activities were not sensitive to the level of sensorimotor noise, which might explain why deterministic simulations can capture the mean walking pattern. However, kinematic and muscle activity variability as well as expected effort were sensitive to the level of sensorimotor noise. Our simulations did not accurately capture the magnitude of experimentally observed variability, probably due to model simplifications (e.g. no neuro-muscular delays). Yet, they captured the structure of the variability well. Stochastic optimal control simulations are a potentially powerful tool to study control of walking. Simulations complement experimental approaches because they enable to test the influence of isolated properties (e.g. magnitude of noise) and to evaluate the underlying control properties (e.g. feedback gains). This is especially relevant to study the origins of balance problems in neurological disorders.

The control policy that emerged from minimising expected effort while stabilising locomotion captured the main features of the experimentally observed structure of the variability. During the step-to-step transition, the variance in joint angles was oriented mostly along the uncontrolled manifold of the centre of mass position. Also the orientation of a line connecting the stance foot to the COM, here called the COM angle, was less variable than the individual joint angles during stance. It has been suggested that a control objective during walking is minimisation of COM kinematic variability, based on the observation that joint angle variability is structured such that COM kinematic variability is small (42,43,46,72). In our simulations, the sole objective was to minimise effort while imposing stability by constraining the state distribution to be periodic (i.e. the variability cannot increase from cycle to cycle) based on a full state feedback controller. The structure of the variability thus emerged and was not induced through penalising COM variability, or through the structure of the feedback controller. In addition, the simulated structure of the variability was not very sensitive to the level of noise. Hence, our simulations suggest that tightly controlling the COM position is the optimal strategy to minimise effort in the presence of noise. Reducing the variability of minimal swing foot clearance during swing might be similarly prioritised. Indeed, the variability of minimal swing foot clearance during swing is not very sensitive to the level of sensorimotor noise and the variability of swing foot clearance seems to be smaller at the moment the foot is closest to the ground than at preceding and following time instants (Fig 6, visual inspection). These results therefore suggest that the structure of movement variability (as observed in experiments) emerges from effort minimisation, and demonstrate how stochastic optimal control simulations can be used to probe the objectives underlying human walking.

The simulated mean kinematics were not sensitive to sensorimotor noise, which might explain why deterministic simulations of locomotion are able to capture mean kinematics and muscle activity in healthy adults (18). This is in contrast with simulations of open-loop arm and eye movements, where accounting for noise was needed to obtain plausible movement patterns (2). Musculoskeletal dynamics might provide a strong incentive towards a specific mean trajectory in gait, as deviations from this mean trajectory could have strong energetic consequences (e.g., due to the interaction between gravity and ground contact, alternative postures such as crouched walking require larger joint moments – particularly knee and hip extension – and therefore greater muscle forces and potentially effort). Hence, we only see deviations in mean kinematics for the highest levels of sensorimotor noise, and these coincide with substantial increases in the expected effort (Fig 4b, Fig F - I in S3 Text). We might have underestimated the sensitivity of the mean walking pattern to sensorimotor noise by neglecting sensorimotor delays and imposing the mean step length. Humans have been observed to reduce mean step length in response to perturbations (73–75). It would be interesting to investigate whether this behaviour would emerge and for which noise levels when imposing only the mean forward speed instead of imposing both step length and step time.

Kinematic variability depends non-monotonically on sensorimotor noise levels, suggesting that the optimal control strategy for minimising effort is not to minimise variability. Importantly, this indicates that the magnitude of the kinematic variability is not necessarily a reflection of the magnitude of the underlying sensorimotor noise. Whereas expected effort increased with sensorimotor noise level, kinematic variance first decreased and then increased with increasing sensory noise level. In the absence of sensory noise, feedback is based on exact information and as it is also instantaneous in our simulations, the optimal strategy is to correct kinematic deviations yielding low variability and expected effort. When there is sensory noise, feedback is based on uncertain information and hence, applying feedback control introduces additional uncertainty. This might explain the high kinematic variance at low sensory noise level (0.001^2^). Tight control might introduce additional uncertainty while such tight control is not needed for stability when the sensory information is accurate. At moderate levels of sensory noise (0.002^2^ - 0.005^2^), kinematic variance was lower than at lower or higher levels of sensorimotor noise indicating that tighter control reduces effort at these sensorimotor noise levels. At higher levels of sensory noise (0.01^2^ - 0.02^2^), both kinematic variance and expected effort increased. This might reflect a reduced ability to limit variability due to less reliable feedback. While we observed different magnitudes of kinematic variance between sensorimotor noise levels, simulations generally underestimated the magnitude of the variance. This might be due to model simplifications, such as the absence of neuromechanical delay that would limit the rate of force development. In our current model, corrective adjustments in excitations immediately result in force. We expect that introducing muscle activation and contraction dynamics and neural delays will result in an overall increase in kinematic and muscle activation variability. Nevertheless, the proposed model captures how variability changes throughout the gait cycle as well as the structure of the variability.

We did not explicitly model different sensory inputs and how their information is combined, yet the proposed framework is a potentially powerful tool to study these processes as well. We consider uncertainty in the state estimation, which is the result of combining delayed, noisy sensory information with prior information (e.g. an internal model). Our approach is not limited to this control structure. It would be insightful to model sensory information from muscles, vision, vestibular, and plantar touch and combine them to generate muscle excitations. The control architecture could be conceptual (e.g. a state estimator and full-state feedback) or inspired by neural pathways (e.g. spinal feedback with limited information but low delay and supraspinal feedback that uses all sensory information but with higher delays). In the current study, we assumed that the feedback error is expressed with respect to the mean states. Our approach does not require this assumption; the reference used for feedback could also be defined as parameters of the control policy and thus be chosen by the optimisation.

Our simulations captured the main features of the structure of observed variability, notwithstanding strong assumptions about the state distribution. In agreement with previous research, we assumed that the distributions of sensorimotor noise and the states of the neuromusculoskeletal dynamics are Gaussian (4,8,37) and we made approximations to propagate this Gaussian state distribution in time. However, it has been shown that the unscented transform is more accurate for nonlinear dynamics than methods based on local linearisation (38). It would be worthwhile to further explore the consequences of the assumptions and approximations made here. In particular, alternative methods that can deal with non-Gaussian distributions have been proposed in literature. For example, the sampling scheme of the unscented transform can be adapted to more accurately propagate the mean and covariance of non-Gaussian distributions (76). Alternatively, the unscented transform can be applied to a Gaussian mixture model, rather than a single Gaussian distribution (77). As these methods are extensions of the unscented transform, they would fit within the proposed approach.

Although we advanced the state of the art, multiple computational challenges remain. A first challenge follows from the size of the optimal control problem (i.e. the number of variables and constraints) and its strong increase with model complexity. Due to the covariance matrix and the increasing number of required sigma points, the number of variables and constraints increases quadratically with the number of states (as opposed to a linear increase in deterministic optimal control problems). This will influence the computational cost of evaluating the nonlinear program (i.e. objective, constraints, and their derivatives) and the cost of finding the direction in which the optimisation variables need to change to advance towards a solution. We have greatly reduced the computational cost of evaluating the nonlinear program (see S2 Text). In our simulations, finding the direction to the next iteration accounted for 93 % of the elapsed real time. This direction is obtained by solving a linear system of equations containing the (derivatives of the) objective and constraints of the nonlinear program. In an optimal control problem, this system of equations is structured because the same dynamics and collocation equations are applied in every mesh interval. By exploiting this structure, the system of equations can be solved 10 times faster (78). Using a structure-exploiting optimal control solver such as FATROP (79) could improve the efficiency of the simulations. However, the convergence of our simulations requires solver features that are currently not implemented in FATROP (e.g. Hessian approximation). Future stochastic optimal control simulations could benefit greatly from the ongoing research into efficient solver algorithms. A second challenge is that the optimal control problem is difficult to solve because it is non-convex. All simulations we ran converged to an optimal solution, but the solver algorithm struggled during most simulations. The primal and dual infeasibilities (which represent the violation of optimality conditions) oscillated without much progress, often for 1000’s of iterations (Fig C in S2 Text). This indicates numerical instability, which is caused by non-convexity and increases with problem size. Hence, the solver might not be able to find a solution when using more complex neuromusculoskeletal models. Therefore, future work should explore modifying the problem formulation, tuning solver parameters, and using different solver algorithms. Alternative simulation approaches, such as reinforcement learning, may avoid these challenges, but will have their own challenges. Some reinforcement learning methods use stochastic algorithms to solve for control parameters (25,32), which is conceptually different from modelling sensorimotor noise. Sensorimotor noise is part of the neuromusculoskeletal model. In the context of reinforcement learning, this means the environment will be stochastic. Describing the noise distribution and the propagation of the state distribution in sufficient detail, while maintaining a reasonable computational cost, will also pose a challenge for reinforcement learning methods. It is currently unclear whether stochastic optimal control, reinforcement learning, or another approach is best suited to simulate walking with complex neuromusculoskeletal models in the presence of uncertainty. Hence, future research should not limit itself to exploring a single method.

In conclusion, we performed stochastic optimal control simulations of walking based on a neuromusculoskeletal model with sensorimotor noise, using a new solution approach. We found that, when minimising expected effort while stabilising locomotion, noise shapes variability but has a small effect on the mean trajectories. Our simulations underestimated the magnitude of the variability, but captured the main features of the experimentally observed structure of the variability. Future work could use our simulations to further probe the neural and musculoskeletal contributions to movement variability in healthy and impaired walking. Simulation code is available at https://codeberg.org/Lars-DHondt/SOC_walking_DHondt2026 under the AGPL v3.0 license. under the AGPL v3.0 license.

## Acknowledgments

We thank Tom Van Wouwe, Wouter Muijres, and Dhruv Gupta for the insightful discussions about stochastic optimal control simulations of human movement.

This work was supported by Research Foundation Flanders (https://www.fwo.be/en/, junior research project fundamental research G0B4222N and G088420N to FDG). The funders had no role in study design, data collection and analysis, decision to publish, or preparation of the manuscript.

## Supporting information

**S1 Text. Additional information about the problem formulation**. (a) smoothed saturation of muscle excitations. (b) derivation of unscented transform parameters. (c) imposing continuity of the state covariance.

**S2 Text. Details about the implementation of the simulation code**. (a) code generation. (b) BLAS. (c) Jacobian helper. (d) Parallel computing. (e) Numerical behaviour of the simulations.

**S3 Text. Additional results**. Kinematics, muscle activations, ground reaction forces, and foot clearance for all simulated noise levels.

## References

1. Faisal AA, Selen LPJ, Wolpert DM. Noise in the nervous system. Nat Rev Neurosci. 2008 Apr;9(4):292–303. doi:10.1038/nrn2258

2. Todorov E. Optimality principles in sensorimotor control. Nat Neurosci. 2004 Sep;7(9):907–15. doi:10.1038/nn1309

3. Harris CM, Wolpert DM. Signal-dependent noise determines motor planning. Nature. 1998 Aug 20;394(6695):780–4. doi:10.1038/29528 PubMed PMID: 9723616.

4. Miyamoto H, Kawato M. Task optimization in the presence of signal-dependent noise (TOPS(α)) model. Int Congr Ser. 2004 Aug 1;1269:105–8. doi:10.1016/j.ics.2004.05.104

5. Todorov E, Jordan MI. Optimal feedback control as a theory of motor coordination. Nat Neurosci. 2002 Nov;5(11):11. doi:10.1038/nn963

6. Cluff T, Crevecoeur F, Scott SH. Tradeoffs in optimal control capture patterns of human sensorimotor control and adaptation [Internet]. bioRxiv; 2019 [cited 2025 Aug 21]. p. 730713. Available from: https://www.biorxiv.org/content/10.1101/730713v1 doi:10.1101/730713

7. Sketch SM, Simpson CS, Crevecoeur F, Okamura AM. Simulating the impact of sensorimotor deficits on reaching performance. In: 2017 International Conference on Rehabilitation Robotics (ICORR) [Internet]. 2017 [cited 2025 Aug 21]. p. 31–7. Available from: https://ieeexplore.ieee.org/abstract/document/8009217 doi:10.1109/ICORR.2017.8009217

8. Haruno M, Wolpert DM. Optimal Control of Redundant Muscles in Step-Tracking Wrist Movements. J Neurophysiol. 2005 Dec;94(6):4244–55. doi:10.1152/jn.00404.2005

9. Van Wouwe T, Ting LH, De Groote F. An approximate stochastic optimal control framework to simulate nonlinear neuro-musculoskeletal models in the presence of noise. PLOS Comput Biol. 2022 Jun 8;18(6):e1009338. doi:10.1371/journal.pcbi.1009338

10. Kuo AD. An optimal state estimation model of sensory integration in human postural balance. J Neural Eng. 2005 Aug;2(3):S235. doi:10.1088/1741-2560/2/3/S07

11. Koelewijn AD, Bogert AJVD. Antagonistic co-contraction can minimize muscular effort in systems with uncertainty. PeerJ. 2022 Apr 7;10:e13085. doi:10.7717/peerj.13085

12. Sinkjaer T, Toft E, Andreassen S, Hornemann BC. Muscle stiffness in human ankle dorsiflexors: intrinsic and reflex components. J Neurophysiol. 1988 Sep;60(3):1110–21. doi:10.1152/jn.1988.60.3.1110

13. Selen LPJ, Beek PJ, van Dieën JH. Impedance is modulated to meet accuracy demands during goal-directed arm movements. Exp Brain Res. 2006 Jun 1;172(1):129–38. doi:10.1007/s00221-005-0320-7

14. Loeb GE, Brown IE, Cheng EJ. A hierarchical foundation for models of sensorimotor control. Exp Brain Res. 1999 Apr 1;126(1):1–18. doi:10.1007/s002210050712

15. Biewener AA, Daley MA. Unsteady locomotion: integrating muscle function with whole body dynamics and neuromuscular control. J Exp Biol. 2007 Sep 1;210(17):2949–60. doi:10.1242/jeb.005801

16. Wagner H, Blickhan R. Stabilizing Function of Skeletal Muscles: an Analytical Investigation. J Theor Biol. 1999 Jul 21;199(2):163–79. doi:10.1006/jtbi.1999.0949

17. Gerritsen KGM, Bogert AJ van den, Hulliger M, Zernicke RF. Intrinsic Muscle Properties Facilitate Locomotor Control—A Computer Simulation Study. Motor Control. 1998 Jul 1;2(3):206–20. doi:10.1123/mcj.2.3.206

18. John CT, Anderson FC, Higginson JS, Delp SL. Stabilisation of walking by intrinsic muscle properties revealed in a three-dimensional muscle-driven simulation. Comput Methods Biomech Biomed Engin. 2013 Apr 1;16(4):451–62. doi:10.1080/10255842.2011.627560 PubMed PMID: 22224406.

19. Haeufle DFB, Grimmer S, Seyfarth A. The role of intrinsic muscle properties for stable hopping—stability is achieved by the force–velocity relation. Bioinspir Biomim. 2010 Feb;5(1):016004. doi:10.1088/1748-3182/5/1/016004

20. Donelan M, Kram R, Kuo A. Mechanical work for step-to-step transitions is a major determinant of metabolic cost of human walking. J Exp Biol. 2003 Jan 1;205:3717–27. doi:10.1242/jeb.205.23.3717

21. Kuo AD, Donelan JM, Ruina A. Energetic Consequences of Walking Like an Inverted Pendulum: Step-to-Step Transitions. Exerc Sport Sci Rev. 2005 Apr;33(2):88.

22. Monaco V, Tropea P, Rinaldi LA, Micera S. Uncontrolled manifold hypothesis: Organization of leg joint variance in humans while walking in a wide range of speeds. Hum Mov Sci. 2018 Feb 1;57:227–35. doi:10.1016/j.humov.2017.08.019

23. Papi E, Rowe PJ, Pomeroy VM. Analysis of gait within the uncontrolled manifold hypothesis: Stabilisation of the centre of mass during gait. J Biomech. 2015 Jan 21;48(2):324–31. doi:10.1016/j.jbiomech.2014.11.024

24. Black DP, Smith BA, Wu J, Ulrich BD. Uncontrolled manifold analysis of segmental angle variability during walking: preadolescents with and without Down syndrome. Exp Brain Res. 2007 Dec 1;183(4):511–21. doi:10.1007/s00221-007-1066-1

25. Qu X. Uncontrolled manifold analysis of gait variability: Effects of load carriage and fatigue. Gait Posture. 2012 Jun 1;36(2):325–9. doi:10.1016/j.gaitpost.2012.03.004

26. Verrel J, Lövdén M, Lindenberger U. Motor-equivalent covariation stabilizes step parameters and center of mass position during treadmill walking. Exp Brain Res. 2010 Nov 1;207(1):13–26. doi:10.1007/s00221-010-2424-y

27. Winter DA. Foot Trajectory in Human Gait: A Precise and Multifactorial Motor Control Task. Phys Ther. 1992 Jan 1;72(1):45–53. doi:10.1093/ptj/72.1.45

28. Afschrift M, Groote FD, Jonkers I. Similar sensorimotor transformations control balance during standing and walking. PLOS Comput Biol. 2021 Jun 25;17(6):e1008369. doi:10.1371/journal.pcbi.1008369

29. Winter DA. Kinematic and kinetic patterns in human gait: Variability and compensating effects. Hum Mov Sci. 1984 Mar 1;3(1):51–76. doi:10.1016/0167-9457(84)90005-8

30. Hicheur H, Terekhov AV, Berthoz A. Intersegmental Coordination During Human Locomotion: Does Planar Covariation of Elevation Angles Reflect Central Constraints? J Neurophysiol. 2006 Sep;96(3):1406–19. doi:10.1152/jn.00289.2006

31. De Groote F, Falisse A. Perspective on musculoskeletal modelling and predictive simulations of human movement to assess the neuromechanics of gait. Proc R Soc B Biol Sci. 2021;288(1946):20202432. doi:10.1098/rspb.2020.2432

32. Denayer M, Alfio E, Díaz MA, Sartori M, De Groote F, De Pauw K, et al. A PRISMA systematic review through time on predictive musculoskeletal simulations. J NeuroEngineering Rehabil. 2025 Jul 4;22(1):149. doi:10.1186/s12984-025-01686-w

33. Falisse A, Serrancoli G, Dembia CL, Gillis J, Jonkers I, De Groote F. Rapid predictive simulations with complex musculoskeletal models suggest that diverse healthy and pathological human gaits can emerge from similar control strategies. J R Soc Interface. 2019;16(157):20190402. doi:10.1098/rsif.2019.0402

34. D’Hondt L, De Groote F, Afschrift M. A dynamic foot model for predictive simulations of human gait reveals causal relations between foot structure and whole-body mechanics. PLOS Comput Biol. 2024 Jun 20;20(6):e1012219. doi:10.1371/journal.pcbi.1012219

35. Vandekerckhove I, D’Hondt L, Gupta D, Van Den Bosch B, Van den Hauwe M, Goemans N, et al. Muscle weakness but also contractures contribute to the progressive gait pathology in children with Duchenne muscular dystrophy: a simulation study. J NeuroEngineering Rehabil. 2025 May 4;22(1):103. doi:10.1186/s12984-025-01631-x

36. Van Den Bosch B, D’Hondt L, Jonkers I, Desloovere K, Van Campenhout A, De Groote F. Estimating the contribution of musculoskeletal impairments to altered gait kinematics in children with cerebral palsy using predictive simulations. J NeuroEngineering Rehabil. 2025 Oct 28;22(1):225. doi:10.1186/s12984-025-01767-w

37. Weng J, Hashemi E, Arami A. Natural Walking With Musculoskeletal Models Using Deep Reinforcement Learning. IEEE Robot Autom Lett. 2021 Apr;6(2):4156–62. doi:10.1109/LRA.2021.3067617

38. Geijtenbeek T. SCONE: Open Source Software for Predictive Simulation of Biological Motion. J Open Source Softw. 2019 Jun 14;4(38):1421. doi:10.21105/joss.01421

39. Song S, Geyer H. Predictive neuromechanical simulations indicate why walking performance declines with ageing. J Physiol. 2018;596(7):1199–210. doi:10.1113/JP275166

40. Song S, Geyer H. A neural circuitry that emphasizes spinal feedback generates diverse behaviours of human locomotion. J Physiol. 2015;593(16):3493–511.

41. Waterval, N. F. J., Veerkamp, K., Geijtenbeek, T., Harlaar, J., Nollet, F., Brehm, M. A., et al. Validation of forward simulations to predict the effects of bilateral plantarflexor weakness on gait. Gait Posture. 2021 Jun 1;87:33–42. doi:10.1016/j.gaitpost.2021.04.020

42. Song S, Geyer H. Evaluation of a neuromechanical walking control model using disturbance experiments. Front Comput Neurosci. 2017;11:15–15. doi:10.3389/fncom.2017.00015

43. Schumacher P, Pierre T, Caggiano V, Kumar V, Schmitt S, Martius G, et al. Emergence of Natural and Robust Bipedal Walking by Learning from Biologically Plausible Objectives. XXX Congress of the International Society of Biomechanics (ISB). 2025.

44. Haeufle DFB, Schmortte B, Geyer H, Müller R, Schmitt S. The Benefit of Combining Neuronal Feedback and Feed-Forward Control for Robustness in Step Down Perturbations of Simulated Human Walking Depends on the Muscle Function. Front Comput Neurosci [Internet]. 2018 [cited 2022 Feb 2];12. Available from: https://www.frontiersin.org/article/10.3389/fncom.2018.00080

45. Zhou B, Zeng H, Wang F, Li Y, Tian H. Efficient and Robust Reinforcement Learning with Uncertainty-based Value Expansion [Internet]. arXiv; 2019 [cited 2025 Sep 1]. Available from: http://arxiv.org/abs/1912.05328 doi:10.48550/arXiv.1912.05328

46. Schumacher P, Häufle D, Büchler D, Schmitt S, Martius G. DEP-RL: Embodied Exploration for Reinforcement Learning in Overactuated and Musculoskeletal Systems [Internet]. arXiv; 2023 [cited 2025 Jul 2]. Available from: http://arxiv.org/abs/2206.00484 doi:10.48550/arXiv.2206.00484

47. Geyer H, Herr H. A Muscle-Reflex Model That Encodes Principles of Legged Mechanics Produces Human Walking Dynamics and Muscle Activities. IEEE Trans Neural Syst Rehabil Eng. 2010;18(3):263–73.

48. Koelewijn AD, van den Bogert AJ. A solution method for predictive simulations in a stochastic environment. J Biomech. 2020 May 7;104:109759. doi:10.1016/j.jbiomech.2020.109759

49. Afschrift M, De Groote F, Verschueren S, Jonkers I. Increased sensory noise and not muscle weakness explains changes in non-stepping postural responses following stance perturbations in healthy elderly. Gait Posture. 2018 Jan 1;59:122–7. doi:10.1016/j.gaitpost.2017.10.003

50. De Groote F. Stochastic optimal control simulations to assess the neuromechanics of human movement. Advances in Motor Learning and Motor Control. 2021 Nov 5.

51. Koelewijn AD. Predictive Simulations of Gait and Their Application in Prosthesis Design [Internet]. Cleveland State University; 2018 [cited 2025 Jul 24]. Available from: https://etd.ohiolink.edu/acprod/odb_etd/etd/r/1501/10?clear=10&p10_accession_num=csu1533901459119777

52. Li W, Todorov E. An Iterative Optimal Control and Estimation Design for Nonlinear Stochastic System. In: Proceedings of the 45th IEEE Conference on Decision and Control [Internet]. 2006 [cited 2025 Dec 1]. p. 3242–7. Available from: https://ieeexplore.ieee.org/abstract/document/4177797 doi:10.1109/CDC.2006.377485

53. Julier SJ, Uhlmann JK. Unscented filtering and nonlinear estimation. Proc IEEE. 2004 Mar;92(3):401–22. doi:10.1109/JPROC.2003.823141

54. Julier SJ, Uhlmann JK, Durrant-Whyte HF. A new approach for filtering nonlinear systems. In: Proceedings of 1995 American Control Conference - ACC’95. 1995. p. 1628–32 vol.3. doi:10.1109/ACC.1995.529783

55. Ozaki N, Campagnola S, Funase R. Tube Stochastic Optimal Control for Nonlinear Constrained Trajectory Optimization Problems. J Guid Control Dyn. 2020 Mar 3;43:1–11. doi:10.2514/1.G004363

56. Ackermann M, van den Bogert AJ. Predictive simulation of gait in rehabilitation. 2010 Annu Int Conf IEEE Eng Med Biol. 2010;5444–7.

57. De Groote F, Pipeleers G, Jonkers I, Demeulenaere B, Patten C, Swevers J, et al. A physiology based inverse dynamic analysis of human gait: potential and perspectives. Comput Methods Biomech Biomed Engin. 2009 Oct 1;12(5):563–74. doi:10.1080/10255840902788587 PubMed PMID: 19319704.

58. De Groote F, Kinney AL, Rao AV, Fregly BJ. Evaluation of Direct Collocation Optimal Control Problem Formulations for Solving the Muscle Redundancy Problem. Ann Biomed Eng. 2016;44(10):2922–36. doi:10.1007/s10439-016-1591-9

59. van den Bogert AJ, Blana D, Heinrich D. Implicit methods for efficient musculoskeletal simulation and optimal control. Procedia IUTAM. 2011 Jan 1;IUTAM Symposium on Human Body Dynamics 2:297–316. doi:10.1016/j.piutam.2011.04.027

60. Delp SL, Anderson FC, Arnold AS, Loan P, Habib A, John CT, et al. OpenSim: Open-Source Software to Create and Analyze Dynamic Simulations of Movement. IEEE Trans Biomed Eng. 2007;54(11):1940–50. doi:10.1109/TBME.2007.901024

61. Seth A, Hicks JL, Uchida TK, Habib A, Dembia CL, Dunne JJ, et al. OpenSim: Simulating musculoskeletal dynamics and neuromuscular control to study human and animal movement. PLOS Comput Biol. 2018 Jul 26;14(7):e1006223. doi:10.1371/journal.pcbi.1006223

62. Falisse A, Serrancolí G, Dembia CL, Gillis J, Groote FD. Algorithmic differentiation improves the computational efficiency of OpenSim-based trajectory optimization of human movement. PLOS ONE. 2019 okt;14(10):e0217730. doi:10.1371/journal.pone.0217730

63. Sherman MA, Seth A, Delp SL. Simbody: multibody dynamics for biomedical research. Procedia IUTAM. 2011 Jan 1;IUTAM Symposium on Human Body Dynamics 2:241–61. doi:10.1016/j.piutam.2011.04.023

64. Zajac FE. Muscle and tendon: properties, models, scaling, and application to biomechanics and motor control. Crit Rev Biomed Eng. 1989;17(4):359–411. PubMed PMID: 2676342.

65. D’Hondt L, Falisse A, Gupta D, Van Den Bosch B, Buurke TJW, Febrer-Nafría M, et al. PredSim: A Framework for Rapid Predictive Simulations of Locomotion. In: 10th IEEE RAS/EMBS BioRob [Internet]. 2024 [cited 2024 Nov 4]. p. 1208–13. Available from: https://ieeexplore.ieee.org/abstract/document/10719735 doi:10.1109/BioRob60516.2024.10719735

66. Anderson FC, Pandy MG. A Dynamic Optimization Solution for Vertical Jumping in Three Dimensions. Comput Methods Biomech Biomed Engin. 1999 Jan 1;2(3):201–31. doi:10.1080/10255849908907988 PubMed PMID: 11264828.

67. Serrancoli G, Falisse A, Dembia C, Vantilt J, Tanghe K, Lefeber D, et al. Subject-Exoskeleton Contact Model Calibration Leads to Accurate Interaction Force Predictions. IEEE Trans Neural Syst Rehabil Eng. 2019;27(8):1597–605.

68. Hunt K, Crossley E. Coefficient of restitution interpreted as damping in vibroimpact. J Appl Mech. 1975.

69. Wolpert DM. Probabilistic models in human sensorimotor control. Hum Mov Sci. 2007 Aug 1;European Workshop on Movement Science 200726(4):511–24. doi:10.1016/j.humov.2007.05.005

70. Ross IM, Karpenko M, Proulx RJ. Path constraints in tychastic and unscented optimal control: Theory, application and experimental results. In: 2016 American Control Conference (ACC) [Internet]. 2016 [cited 2025 Nov 19]. p. 2918–23. Available from: https://ieeexplore.ieee.org/abstract/document/7525362 doi:10.1109/ACC.2016.7525362

71. Ross IM, Proulx RJ, Karpenko M. Unscented optimal control for space flight. In: 24th International Symposium on Space Flight Dynamics (ISSFD) [Internet]. 2014 [cited 2025 Mar 20]. p. 1–12. Available from: https://www.issfd.org/ISSFD_2014/ISSFD24_Paper_S12-5_Karpenko.pdf

72. Andersson JAE, Gillis J, Horn G, Rawlings JB, Diehl M. CasADi: a software framework for nonlinear optimization and optimal control.Math Program Comput. 2019 Mar 1;11(1):1–36. doi:10.1007/s12532-018-0139-4

73. Wächter A, Biegler LT. On the implementation of an interior-point filter line-search algorithm for large-scale nonlinear programming. Math Program. 2006 Mar 1;106(1):25–57. doi:10.1007/s10107-004-0559-y

74. Hogg JD, Scott JA. HSL_MA97 : a bit-compatible multifrontal code for sparse symmetric systems. Rutherford Appleton Lab Tech Rep [Internet]. 2011 [cited 2025 Mar 6];RAL-TR-2011-024. Available from: https://epubs.stfc.ac.uk/work/61445

75. HSL. A collection of Fortran codes for large scale scientific computation. [Internet]. [cited 2026 Feb 8]. Available from: https://www.hsl.rl.ac.uk/

76. Betts JT. Practical Methods for Optimal Control and Estimation Using Nonlinear Programming, Second Edition [Internet]. Society for Industrial and Applied Mathematics; 2010 [cited 2025 Oct 10]. 442 p. (Advances in Design and Control). Available from: https://epubs.siam.org/doi/book/10.1137/1.9780898718577 doi:10.1137/1.9780898718577

77. Lawson CL, Hanson RJ, Kincaid DR, Krogh FT. Basic Linear Algebra Subprograms for Fortran Usage. ACM Trans Math Softw. 1979 Sep 1;5(3):308–23. doi:10.1145/355841.355847

78. AOCL-BLIS [Internet]. AMD; [cited 2025 Nov 25]. Available from: https://www.amd.com/en/developer/aocl/blis.html

79. Afschrift M, Pitto L, Aerts W, van Deursen R, Jonkers I, De Groote F. Modulation of gluteus medius activity reflects the potential of the muscle to meet the mechanical demands during perturbed walking. Sci Rep. 2018 Aug 3;8(1):11675. doi:10.1038/s41598-018-30139-9

80. Falisse A, Afschrift M, Groote FD. Modeling toes contributes to realistic stance knee mechanics in three-dimensional predictive simulations of walking. PLOS ONE. 2022 Jan 25;17(1):e0256311. doi:10.1371/journal.pone.0256311

81. Nene A, Byrne C, Hermens H. Is rectus femoris really a part of quadriceps?: Assessment of rectus femoris function during gait in able-bodied adults. Gait Posture. 2004 Aug 1;20(1):1–13. doi:10.1016/S0966-6362(03)00074-2

82. Dingwell J, John J, Cusumano J. Do Humans Optimally Exploit Redundancy to Control Step Variability in Walking? PLoS Comput Biol. 2010 Jul 15;6:e1000856. doi:10.1371/journal.pcbi.1000856

83. Voloshina AS, Kuo AD, Daley MA, Ferris DP. Biomechanics and energetics of walking on uneven terrain. J Exp Biol. 2013 Nov 1;216(21):3963–70. doi:10.1242/jeb.081711

84. O’Connor SM, Kuo AD. Direction-Dependent Control of Balance During Walking and Standing. J Neurophysiol. 2009 Sep;102(3):1411–9. doi:10.1152/jn.00131.2009 PubMed PMID: 19553493; PubMed Central PMCID: PMC2746770.

85. Muijres W, Afschrift M, Ronsse R, Groote FD. Speeding up, not slowing down, decreases the energy needed to control walking balance [Internet]. bioRxiv; 2025 [cited 2026 Feb 27]. p. 2025.03.25.645241. Available from: https://www.biorxiv.org/content/10.1101/2025.03.25.645241v1 doi:10.1101/2025.03.25.645241

86. Ebeigbe D, Berry T, Whalen AJ, Norton MM, Simon D, Sauer TD, et al. Generalized unscented transformation for forecasting non-Gaussian processes. Phys Rev E. 2025 May 27;111(5):054135. doi:10.1103/PhysRevE.111.054135

87. Luo X, Moroz IM, Hoteit I. Scaled unscented transform Gaussian sum filter: Theory and application. Phys Nonlinear Phenom. 2010 May 15;239(10):684–701. doi:10.1016/j.physd.2010.01.022

88. Vanroye L, De Schutter J, Decré W. A generalization of the Riccati recursion for equality-constrained linear quadratic optimal control. Optim Control Appl Methods. 2024;45(1):436–54. doi:10.1002/oca.3064

89. Vanroye L, Sathya A, De Schutter J, Decré W. FATROP: A Fast Constrained Optimal Control Problem Solver for Robot Trajectory Optimization and Control. In: 2023 IEEE/RSJ International Conference on Intelligent Robots and Systems (IROS) [Internet]. 2023 [cited 2026 Feb 28]. p. 10036–43. Available from: https://ieeexplore.ieee.org/abstract/document/10342336 doi:10.1109/IROS55552.2023.10342336

90. Kidziński Ł, Mohanty SP, Ong C, Huang Z, Zhou S, Pechenko A, et al. Learning to Run challenge solutions: Adapting reinforcement learning methods for neuromusculoskeletal environments [Internet]. arXiv; 2018 [cited 2026 Mar 2]. Available from: http://arxiv.org/abs/1804.00361 doi:10.48550/arXiv.1804.00361

