## Supplementary material for "Stochastic optimal control simulations of walking: potential and perspective": S1 Text

### S1 Text. Problem formulation

**Table A. Summary of symbols and notations, repeated from main text.**

| Symbol or notation | Meaning |
|---|---|
| $x$ | State vector |
| $\boldsymbol{P}$ | State covariance matrix |
| $\boldsymbol{L}$ | Cholesky factor of state covariance matrix ($\boldsymbol{P} = \boldsymbol{L}\boldsymbol{L}^T$) |
| $u$ | Control vector |
| $e_{ff}$ | Feedforward excitation |
| $\boldsymbol{K}_{fb}$ | Feedback gains |
| $e_{tot}$ | Total excitation |
| $\xi$ | Noise vector |
| $\boldsymbol{\Xi}$ | Noise covariance matrix |
| $\Xi_{\mathrm{m}}$ $\Xi_{\mathrm{s}}$ | Noise level (motor resp. sensory) |
| $c$ | Scalar determining the spread of the sigma points (here, 3) |
| ${}^{m}W$ | Vector with weights to calculate mean of sigma points |
| ${}^{c}\boldsymbol{W}$ | Diagonal matrix with weights to calculate covariance of sigma points |
| $\dot{x}$ | Derivative to time |
| $\bar{x}$ | Mean |
| ${}^{a}x$ | Augmented state, including both state and noise variables |
| $N_m$ | The number of mesh intervals (here, 64) |
| $N_c$ | The order of the Radau collocation scheme (here, 3) |
| $N_\sigma$ | The number of sigma points (here, 109) |
| $x_k$ | Variable at k^th^ mesh point or interval |
| $x_{k,j}$ | Variable at j^th^ collocation point of k^th^ mesh interval |
| $x^{(i)}$ | Variable at i^th^ sigma point |
| $x_{k\|k-1}$ | Variable calculated based on previous mesh point or interval |
| $x + \boldsymbol{L}$ | The vector $x$ is added to every column of the matrix $\boldsymbol{L}$ |

### Smoothed saturation

Total excitation cannot go below zero or exceed maximal excitation. In other words, if excitation is at its lowest level, inhibitory feedback will not reduce it further, and if excitation is already maximal, additional excitatory drive will not lead to increased activation of the muscle. We modelled this saturation effect as a piece-wise function of a hyperbola, a linear part, and a hyperbola (Fig A). This function increases monotonically and is $C^1$ continuous, making it suitable for gradient-based optimisation. The implementation is available on https://codeberg.org/Lars-DHondt/SOC_walking_DHondt2026/src/branch/main/Functions/smoothed_saturation.m.

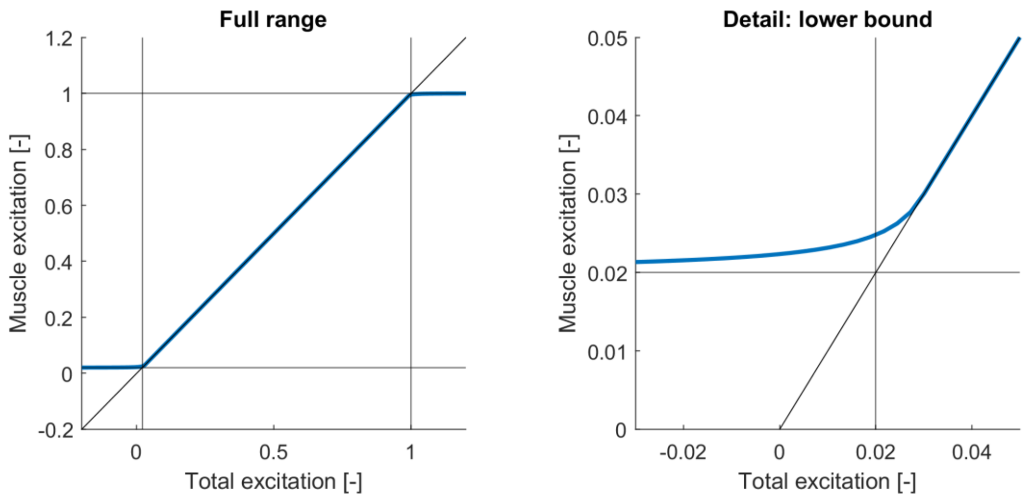


**Fig A. Smoothed saturation.**
Saturation of muscle excitations is approximated as a piece-wise function of a hyperbola, a linear part, and a hyperbola. This function is monotonically increasing and $C^1$ continuous.

### Unscented transform parameters

A distribution of $N_x$ augmented states (i.e. number of states and noise sources) can be represented by $2N_x + 1$ sigma points: the mean point and $2N_x$ points on the c$^{th}$ covariance contour (1). The value of c and the weights used to calculate the mean and covariance of the sigma points have to be chosen such that the original mean and covariance are reconstructed exactly. These criteria do not uniquely determine the parameter values, allowing for additional tuning. The value of c and the weights on the other points ($W^{(i)}$) can be expressed in function of the weight on the mean point ($W^{(0)}$) (1)

$$c = \sqrt{\frac{N_x}{1 - W^{(0)}}} \tag{1}$$

$$W^{(i)} = \frac{1 - W^{(0)}}{2N_x} \tag{2}$$

There are different options for $W^{(0)}$, which we first explain followed by a motivation of our choice. The weight on the mean point is often chosen to be *0* or according to

$$W^{(0)} = 1 - N_x/3\,. \tag{3}$$

When calculating the covariance, the mean point can be given additional weight to improve the accuracy of the transform by incorporating information from the diagonal elements of the kurtosis tensor

$$^{c}W^{(0)} = W^{(0)} + \beta \tag{4}$$

where $\beta = 2$ for a Gaussian distribution (1).

When the number of augmented states increases, equation 3 leads to negative weight values. While choosing a negative value is allowed (1), for optimal control it leads to undesired behaviour. Consider the expected effort (i.e. activation squared) of a single muscle

$$\mathbb{E}(a^2) = W^{(0)}\left(a^{(0)}\right)^2 + W^{(i)}\left(a^{(1)}\right)^2 + \cdots + W^{(i)}\left(a^{(2N_x)}\right)^2 \tag{5}$$

If $W^{(0)}$ is negative, the effort at the mean point would contribute to the expected effort with a negative weight. Thus the lowest expected effort can be achieved by maximising the effort of the mean point and minimising the effort of the other points. We observed that using such nonconvex cost function resulted in difficulties for the optimisation solver. In the cases where the simulations did converge to a solution, the sigma points with negative (or zero) weight had much higher activations than the others.

Therefore, we opted to modify the sampling scheme of the unscented transform such that each point contributes equally to the mean

$$^{m}W^{(0)} = {^{m}W^{(i)}} = \frac{1}{2N_x + 1} \tag{6}$$

Substituting eq. 6 in eq. 1 yields

$$c = \sqrt{N_x + 1/2} \tag{7}$$

For the neuromusculoskeletal model considered in this paper, which has 18 states and 36 noise sources, $c = 7.38$. However, sampling that far from the mean might not yield good approximations due to non-linearities (dynamics around the mean is different than dynamics far away from the mean) and might even lead to unphysiological states (e.g. joint angles outside range of motion. The spread of the sigma points can be controlled by employing the scaled unscented transform (2) but this method yields a negative weight for the mean point, which is undesirable as explained above. We therefore opted to decouple the weights used for the mean and covariance of the sigma points. The weights need to be chosen such that the sigma points encode the mean and covariance of the initial distribution (1). Consider the sigma point set

$$\boldsymbol{\chi} = [\bar{x} \quad \bar{x} + c\sqrt{\boldsymbol{P}} \quad \bar{x} - c\sqrt{\boldsymbol{P}}\,] \tag{8}$$

The covariance of the sigma points is given by

$$\boldsymbol{P} = (\boldsymbol{\chi} - \bar{x})\,{}^{\boldsymbol{c}}\boldsymbol{W}\,(\boldsymbol{\chi} - \bar{x})^T \tag{9}$$

Since ${}^{\boldsymbol{c}}\boldsymbol{W} \equiv diag\left({}^{c}W^{(0)}, {}^{c}W^{(i)}, \dots, {}^{c}W^{(i)}\right)$, the equation can be rewritten as

$$\boldsymbol{P} = {}^{c}W_0\, 0_{N_x x1} + \left(c\sqrt{\boldsymbol{P}}\right) {}^{c}W^{(i)} \left(c\sqrt{\boldsymbol{P}}\right)^T + \left(-c\sqrt{\boldsymbol{P}}\right) {}^{c}W^{(i)} \left(-c\sqrt{\boldsymbol{P}}\right)^T$$

$$\Leftrightarrow \boldsymbol{P} = 2c\,{}^{c}W^{(i)}\left(\sqrt{\boldsymbol{P}}\right)\left(\sqrt{\boldsymbol{P}}\right)^T \tag{10}$$

This condition is satisfied if

$${}^{c}W^{(i)} = \frac{1}{2c^2}. \tag{11}$$

where *c* is a parameter that can be chosen. The covariance correction term can still be included:

$${}^{c}W^{(0)} = \frac{1}{2c^2} + \beta. \tag{12}$$

Here, we used *c = 3* in analogy with chance-constraints being placed on the 3rd covariance contour. Because we assume the state distribution to be Gaussian, we set $\beta = 2$.

### Covariance constraints

We evaluated the computational cost of multiple methods to impose the continuity of the state covariance matrix between two adjacent mesh intervals (equation 14 in main text) and selected the approach with the lowest cost. To improve the readability, we introduce the matrix **Y** that expresses the states at the sigma points relative to the mean state

$$\boldsymbol{Y} \equiv \boldsymbol{\chi}_{\boldsymbol{k+1|k}} - \bar{x}_{k+1} \tag{13}$$

and omit the subscript k+1 (all variables in this section belong to mesh point k+1). The first way to express the continuity is an implicit formulation using the covariance matrices

$$\boldsymbol{\varepsilon} = \boldsymbol{L}\boldsymbol{L}^{\boldsymbol{T}} - \boldsymbol{Y}\,{}^{c}\boldsymbol{W}\,\boldsymbol{Y}^{T} \tag{14}$$

The second way, is to impose continuity on the Cholesky factor of the covariance matrices.

$$\boldsymbol{\varepsilon} = \boldsymbol{L} - chol(\boldsymbol{Y}\,{}^{c}\boldsymbol{W}\,\boldsymbol{Y}^{T}) \tag{15}$$

This is most comparable to explicit propagation of the covariance matrix as is typically done when using unscented transform for filtering, estimation (1), or optimal control (3). The third way is inspired by the square-root unscented Kalman filter (4). Rather than computing the Cholesky decomposition of the product of a matrix with its transpose, the Cholesky factor can be computed directly through a QR decomposition. In general, any real n×m matrix **A** (with m ≥ n) can be decomposed into the product of an m×n matrix with orthogonal columns (**Q**) and an n×n upper triangular matrix (**R**):

$$\boldsymbol{A}^{\boldsymbol{T}} = \boldsymbol{Q}\boldsymbol{R} \tag{16}$$

If **A** has full row rank, it follows that

$$chol\left(\boldsymbol{A}\boldsymbol{A}^{\boldsymbol{T}}\right) = \boldsymbol{R}^{\boldsymbol{T}} \tag{17}$$

(see Theorem 5.2.2 and proof in (5)). Here, we define the matrix **A** as

$$\boldsymbol{A} \equiv \boldsymbol{Y}\,{}^{c}\boldsymbol{W}^{1/2} \tag{18}$$

Combining equations 15 – 18 then yields an expression to impose continuity on the Cholesky factor of the covariance matrices

$${}^{c}\boldsymbol{W}^{1/2}\boldsymbol{Y}^{\boldsymbol{T}} = \boldsymbol{Q}\boldsymbol{R} \tag{19}$$

$$\boldsymbol{\varepsilon} = \boldsymbol{L} - \boldsymbol{R}^{T} \tag{20}$$

The constraint violation $\boldsymbol{\varepsilon}$ is symmetric or lower triangular by definition. Either way, only its lower triangular elements should be imposed as constraints. The Cholesky and QR factorisation can be performed symbolically using CasADi's build-in functions (6). Hence, exact derivatives can be obtained for each constraint formulation approach. Comparing the computational cost of evaluating the constraint and its Jacobian, it is clear that the implicit formulation is the most efficient (Fig B).

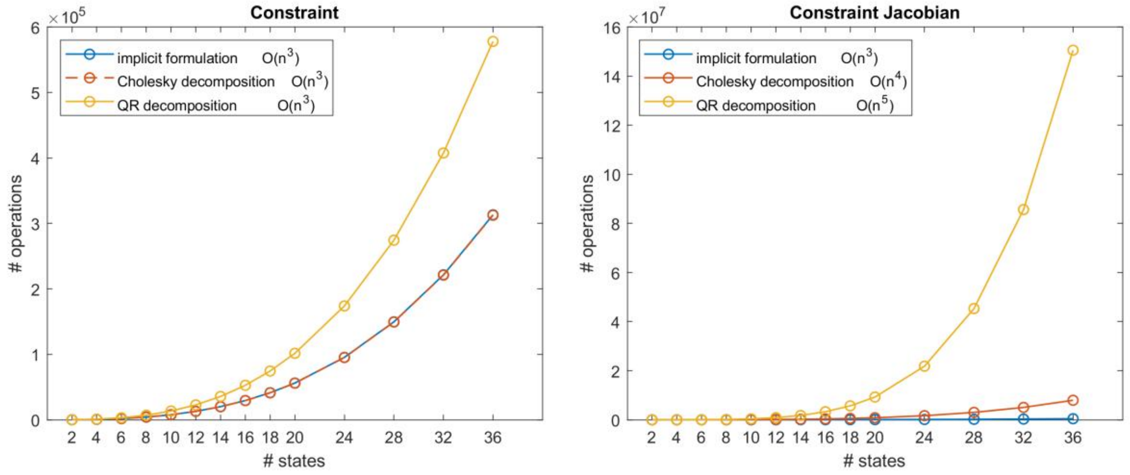


**Fig B. Computational cost of evaluating covariance continuity constraints.**
Cost is expressed as the number of scalar operations (e.g. multiplying two scalars or taking the square root of a scaler) in the expression graph.

### References


1. Julier SJ, Uhlmann JK. Unscented filtering and nonlinear estimation. Proc IEEE. 2004 Mar;92(3):401–22. doi:10.1109/JPROC.2003.823141

2. Julier SJ. The scaled unscented transformation. In: Proceedings of the 2002 American Control Conference (IEEE Cat. No.CH37301). 2002. p. 4555–9 vol.6. doi:10.1109/ACC.2002.1025369

3. Ozaki N, Campagnola S, Funase R. Tube Stochastic Optimal Control for Nonlinear Constrained Trajectory Optimization Problems. J Guid Control Dyn. 2020 Mar 3;43:1–11. doi:10.2514/1.G004363

4. Van der Merwe R, Wan EA. The square-root unscented Kalman filter for state and parameter-estimation. In: 2001 IEEE International Conference on Acoustics, Speech, and Signal Processing. Proceedings (Cat. No.01CH37221). 2001. p. 3461–4 vol.6. doi:10.1109/ICASSP.2001.940586

5. Golub GH, Loan CFV. Matrix Computations. JHU Press; 1996. 734 p.

6. Andersson JAE, Gillis J, Horn G, Rawlings JB, Diehl M. CasADi: a software framework for nonlinear optimization and optimal control. Math Program Comput. 2019 Mar 1;11(1):1–36. doi:10.1007/s12532-018-0139-4
