## Supplementary material for "Stochastic optimal control simulations of walking: potential and perspective": S2 Text

### S2 Text. Simulation code implementation

**Table A. Summary of symbols and notations, repeated from main text.**

| Symbol or notation | Meaning |
|---|---|
| $x$ | State vector |
| $\boldsymbol{P}$ | State covariance matrix |
| $\boldsymbol{L}$ | Cholesky factor of state covariance matrix ($\boldsymbol{P} = \boldsymbol{L}\boldsymbol{L}^T$) |
| $u$ | Control vector |
| $e_{ff}$ | Feedforward excitation |
| $\boldsymbol{K}_{fb}$ | Feedback gains |
| $e_{tot}$ | Total excitation |
| $\xi$ | Noise vector |
| $\boldsymbol{\Xi}$ | Noise covariance matrix |
| $\Xi_m$ $\Xi_s$ | Noise level (motor resp. sensory) |
| $c$ | Scalar determining the spread of the sigma points (here, 3) |
| $^mW$ | Vector with weights to calculate mean of sigma points |
| $^c\boldsymbol{W}$ | Diagonal matrix with weights to calculate covariance of sigma points |
| $\dot{x}$ | Derivative to time |
| $\bar{x}$ | Mean |
| $^ax$ | Augmented state, including both state and noise variables |
| $N_m$ | The number of mesh intervals (here, 64) |
| $N_c$ | The order of the Radau collocation scheme (here, 3) |
| $N_\sigma$ | The number of sigma points (here, 109) |
| $x_k$ | Variable at k$^{th}$ mesh point or interval |
| $x_{k,j}$ | Variable at j$^{th}$ collocation point of k$^{th}$ mesh interval |
| $x^{(i)}$ | Variable at i$^{th}$ sigma point |
| $x_{k\|k-1}$ | Variable calculated based on previous mesh point or interval |
| $x + \boldsymbol{L}$ | The vector $x$ is added to every column of the matrix $\boldsymbol{L}$ |

### Code generation

To reduce the computational cost, most calculations were offloaded to compiled code instead of evaluating them in CasADi's built-in virtual machine. We implemented the dynamics in MATLAB, then used CasADi's built-in code generator (1) to create C code with functions to evaluate the path constraints and their Jacobian. The reduction in evaluation time that can be achieved with compiled code is due to the code optimisation performed by the compiler. Therefore, we tested different compilers and code optimisation levels. We compared the C compilers from three different software development toolkits available for Windows: Microsoft Visual Studio (Community edition 2022), Intel oneAPI (version 2024.0), and MSYS2 (GNU C compiler version 14.2.0). We tested four different levels of code optimisation: no optimisation (Od/O0), moderate (O2), aggressive (O3), aggressive and explicitly state that the processor supports the AVX512 instruction set (O3+AVX512). The exact behaviour of each level may vary between compilers. Since the C compiler in Microsoft Visual Studio does not include aggressive code optimisation, only two levels were used there. Support for the AVX512 instruction set was specified by passing */QxCOMMON-AVX512* (intel) or *-mprefer-vector-width=512* (GNU) to the compiler. All tests were performed on two computers running Windows 11: a laptop with an *Intel Core i7-11850H* CPU (8 cores, 2.50 GHz), and a desktop with an *AMD Ryzen Threadripper Pro 7975WX* CPU (32 cores, 4 GHz). All compiler test cases used the same source file. This file contained C code to evaluate the dynamics of a preliminary neuromusculoskeletal model (with similar complexity as the final model) and the Jacobian of the dynamics at 8 points.

The intel compiler resulted in the lowest evaluation time, both on intel and AMD hardware (Fig **A**). On the desktop computer (which was used to run the full simulations) evaluating the dynamics function was fastest when using aggressive code optimisation and targeting AVX512 instructions. Increasing the code optimisation level beyond moderate (O2) had little effect on duration of Jacobian evaluations. Hence, we selected the intel compiler with aggressive code optimisation and targeting AVX512 instructions for use in the simulations.

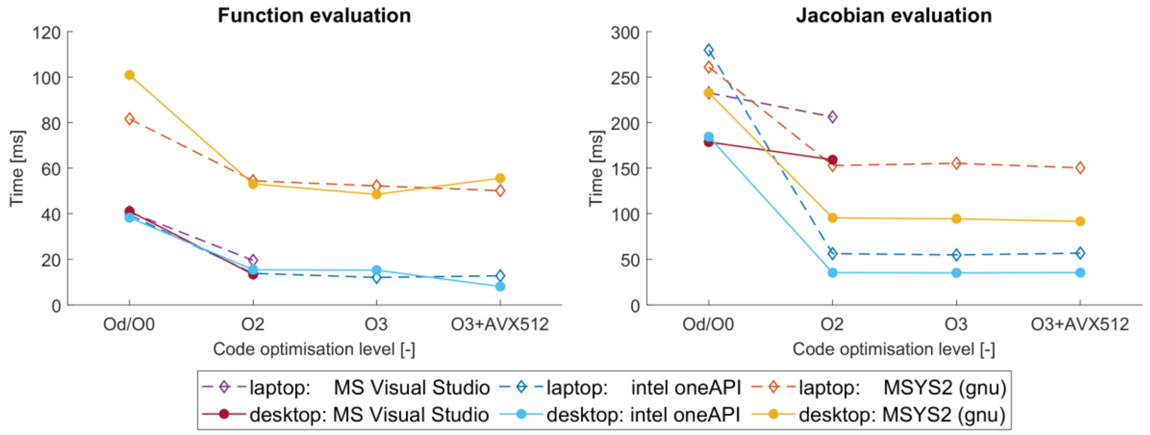


**Fig A. Comparison of compilers and code optimisation levels.**
Time of each evaluation was calculated as the average duration over 10000 repetitions.

### Basic Linear Algebra Subprograms

The path constraints expressing the continuity of the state covariance (equation 14 in main text) consist of multiplications and additions with dense matrices. The computational cost of evaluating these matrix computations scales with the size of the matrices (i.e. the number of states) to the third power, thus, could pose a bottleneck for using more complex models. To speed up the evaluation, we relied on a Basic Linear Algebra Subprograms (BLAS) (2) library that was optimised for our hardware (3).

The constraint (equation 12 in main text) has to be formulated as a combination of BLAS routines. To improve the readability, we introduce

$$\boldsymbol{Y} \equiv \boldsymbol{\chi_{k+1|k}} - \bar{x}_{k+1} \quad (1)$$

and omit the subscript k+1. Then, equation 12 (main text) becomes

$$\boldsymbol{\varepsilon} = \boldsymbol{LL^T} - \boldsymbol{Y}\,{}^c\boldsymbol{W}\,\boldsymbol{Y}^T \quad (2)$$

where the weights are known constants (see S1 Text)

$${}^c\boldsymbol{W} \equiv diag\left({}^cW^{(i)} + \beta, {}^cW^{(i)}, \dots, {}^cW^{(i)}\right) \quad (3)$$

Equation 2 can be written as a sequence of symmetric rank-k updates. These exploit the structure of the matrices and are therefore more efficient than general matrix-matrix multiplications (4). The general form of a symmetric rank-k update is

$$\boldsymbol{C} \leftarrow \boldsymbol{C} + \alpha\boldsymbol{AA^T} \quad (4)$$

where $\alpha$ is a scalar, **C** is a symmetric $n\times n$ matrix, and **A** is an $n\times k$ matrix. Substituting equation 3 into equation 2, and denoting the first column of **Y** as $y^{(0)}$, gives

$$\boldsymbol{\varepsilon} = \boldsymbol{LL^T} - \left({}^cW^{(i)}\boldsymbol{YY^T} + \beta y^{(0)}\left(y^{(0)}\right)^T\right) \quad (5)$$

This is a sequence of three symmetric rank-k updates, where the third is a rank-1 update, and can be implemented with BLAS routines. Because **ε** is symmetric by definition, only its lower triangular elements are imposed as constraints.

Since the constraint was implemented in BLAS and not with CasADi's symbolic variables, its derivative cannot be obtained with algorithmic differentiation. Therefore, we also implemented a function to evaluate its (exact) derivative. The Jacobian of the result of a symmetric rank-k update (equation 4) to the elements of **A**, and therefore also the Jacobian of the constraint (equation 5) to the elements of **L** and **Y**, can be obtained as follows: for illustration, consider **A** as a 3×2 matrix:

$$\boldsymbol{A} \equiv \begin{bmatrix} a_{11} & a_{12} \\ a_{21} & a_{22} \\ a_{31} & a_{32} \end{bmatrix} \quad (6)$$

The update term in equation 4 is then

$$\alpha\boldsymbol{AA^T} = \underbrace{\alpha\begin{bmatrix} a_{11}^2 & a_{11}a_{21} & a_{11}a_{31} \\ a_{11}a_{21} & a_{21}^2 & a_{11}a_{31} \\ a_{11}a_{31} & a_{11}a_{31} & a_{31}^2 \end{bmatrix}}_{1st\ column\ of\ \boldsymbol{A}} + \underbrace{\alpha\begin{bmatrix} a_{12}^2 & a_{12}a_{22} & a_{12}a_{32} \\ a_{12}a_{22} & a_{22}^2 & a_{12}a_{32} \\ a_{12}a_{32} & a_{12}a_{32} & a_{32}^2 \end{bmatrix}}_{2nd\ column\ of\ \boldsymbol{A}} \quad (7)$$

This symmetric matrix can be described by only its lower triangular elements, i.e. the matrix $\boldsymbol{\mathcal{A}}$. The Jacobian of this matrix is

$$\frac{\partial\boldsymbol{\mathcal{A}}}{\partial\boldsymbol{A}} = \alpha\begin{bmatrix} 2a_{11} & 0 & 0 & 2a_{12} & 0 & 0 \\ a_{21} & a_{11} & 0 & a_{22} & a_{12} & 0 \\ a_{31} & 0 & a_{11} & a_{32} & 0 & a_{12} \\ 0 & 2a_{21} & 0 & 0 & 2a_{22} & 0 \\ 0 & a_{31} & a_{21} & 0 & a_{32} & a_{22} \\ \underbrace{0 \quad 0 \quad a_{31}}_{1st\ column\ of\ \boldsymbol{A}} & & & \underbrace{0 \quad 0 \quad a_{32}}_{2nd\ column\ of\ \boldsymbol{A}} & & \end{bmatrix} \quad (8)$$

This Jacobian matrix consists of the elements of **A**, scaled by a factor $\alpha$ or $2\alpha$, and arranged according to a pattern. Both the scale factors and patterns are known before starting the simulation. To evaluate the Jacobian matrix corresponding to equation 5, the Jacobian matrices with respect to each column of $\boldsymbol{L}$ and $\boldsymbol{Y}$ are evaluated and then concatenated horizontally.

We compared our implementation against the reference implementation using CasADi symbolics that ran in CasADi's virtual machine (i.e. the default option) (1). We also used CasADi to generate C code with functions to evaluate the constraint and its Jacobian, and compiled these into a library (i.e. the conventional method to speed up evaluations). The BLAS implementation was fastest for evaluating the constraints, both the BLAS and compiled library were fastest to evaluate the constraint Jacobian (Table B). The latter is likely because the evaluation of the Jacobian requires few calculation and hence benefits little from using BLAS. The BLAS implementation was overall fastest, so we used that in the simulations.

The source code (*LLT-YWYT.c*) and a MATLAB function to perform just-in-time compilation (*jitLLT_YWYT.m*) are available on https://codeberg.org/Lars-DHondt/SOC_walking_DHondt2026/src/branch/main/codegen.

**Table B. Benchmark of state covariance continuity constraint implementation.**
Time to evaluate the constraint and its Jacobian (averaged over 30000 repetitions).

| | Constraint [μs] | Jacobian [μs] |
|---|---|---|
| Virtual Machine | 54 | 195 |
| Compiled Library | 14 | 71 |
| BLAS | 6 | 73 |

### Jacobian helper

When evaluating the constraint Jacobian, most of the computational cost comes from evaluating the Jacobian of the constraints corresponding to the sigma points (i.e. dynamics and collocation equations). Typically, the constraint function is first repeated over all sigma points (or mesh points in general optimal control problems). Then, the Jacobian of all these constraints with respect to all inputs is used to construct the constraint Jacobian of the nonlinear program. Here, we explored an alternative approach: the Jacobian expression is first derived for the constraints of a single sigma point. Then, this Jacobian expression is repeated over all sigma points. We observed that evaluating the latter was around 25 times faster than evaluating the former.

The alternative approach does result in a different structure of the resulting Jacobian matrix: the blocks corresponding to the sigma points are concatenated horizontally instead of diagonally. These matrices are usually sparce, so CasADi stores them using the compressed column format (1). In this format, a sparse matrix is represented as a vector containing the values of the structural nonzero elements and a vector encoding the position of each value in the sparse matrix. The first vector, whose values are updated at every evaluation, is identical between diagonally or horizontally concatenated blocks. Only the second vector, which is set at initialisation, is different.

We created a helper file that, when combined with generated source code to evaluate the repeated Jacobian expression, can be compiled into a library that returns the Jacobian of the repeated function. This helper file includes a function that adapts the vector encoding the positions of the structural non-zeros to represent a block-diagonal matrix, which is run once at initialisation.

To test the effect of this helper function, we compared the elapsed time of evaluating the constraints on implicit dynamics and collocation equation, and of evaluating their Jacobian, for all 109x64 sigma points. We tested the combined effect of the Jacobian helper and using a library compiled with code optimisation, since their implementations are linked. We observed that this combination provided a considerable speed-up over using CasADi's built-in virtual machine (Table C). Evaluating the constraints was 20 times faster, evaluating their Jacobian was over 450 times faster.

The source code (*jac_map_fun.c*) and a MATLAB function to perform just-in-time compilation (*jitMapFun.m*) are available on https://codeberg.org/Lars-DHondt/SOC_walking_DHondt2026/src/branch/main/codegen.

**Table C. Benchmark of Jacobian implementation.**
Time to evaluate the constraints on implicit dynamics and collocation equation, and their Jacobian, for all 109x64 sigma points. Durations were averaged over 10 repetitions.

| | Constraint [ms] | Jacobian [ms] |
|---|---|---|
| Virtual Machine | 319 | 26643 |
| Compiled library with Jacobian helper | 16 | 58 |

### Parallel computing

We used parallel computing to efficiently use computational resources. The computational efficiency of simulations can benefit from parallelism on three levels: First, we exploited instruction-level parallelism (i.e. modern processors can do element-wise addition or multiplication of two small vectors instead of only scalars) by using aggressive code optimisation and preferring AVX512 instructions (vector length 8), and using a BLAS library that is optimised for our specific hardware (3). The optimisation solvers also use BLAS (5–7). Second, we exploited task parallelism to distribute the evaluation of constraints and their Jacobian at the different sigma points over multiple CPU cores (multi-threading via openmp). The linear solver also relies on openmp to parallelise its calculations (6). Third, we used the Parallel Computing Toolbox in MATLAB (The Mathworks Inc., USA) to run multiple simulations at the same time.

To select the number of CPU threads assigned to function evaluations and to the linear solver, we tested their effect on the simulation time. In these tests, we ran simulations (200 iterations) with a preliminary model. The time spent on evaluating the functions that describe the nonlinear program (objective, constraints, and their derivatives) was hardly influenced by multithreading (**Fig B**a). The time spent in the solver (i.e. ipopt (5) and ma97 (6,7)) was lowest when assigning 16 threads to each instance of ma97 (**Fig B**b). Given the 32-core CPU with hyperthreading, running four simulations in parallel was expected to use all 64 virtual CPU cores.

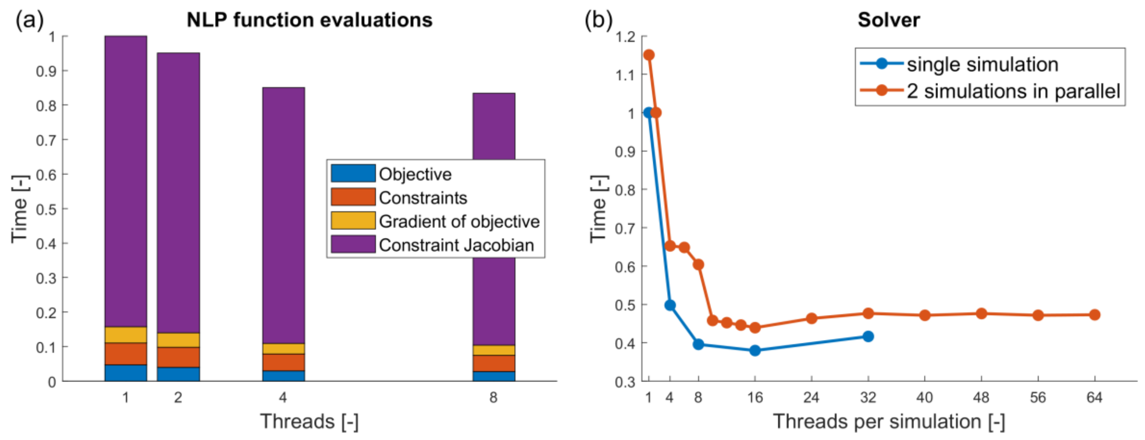


**Fig B. The effect of multithreading on the duration of evaluating the NLP functions.**

### Numerical behaviour of the simulations

All simulations we ran converged to an optimal solution. Evaluation of functions (e.g. constraints) accounted for only 5% of the time (Table D), due to their computationally efficient implementation. We did observe that the solver algorithm struggled during most simulations. The primal and dual infeasibilities (which represent the violation of optimality conditions) oscillated without much progress, often for 1000's of iterations (Fig C). This indicates numerical instability, which is caused by non-convexity and increases with problem size.

**Table D. Contribution of function evaluations to total elapsed real time.**

| Function | Time (mean) [%] | Time (range) [%] |
|---|---|---|
| Objective | 0.13 | 0.083 – 0.195 |
| Constraints | 0.12 | 0.079 – 0.191 |
| Gradient of objective | 0.14 | 0.103 – 0.227 |
| Constraint Jacobian | 4.74 | 3.594 – 5.337 |

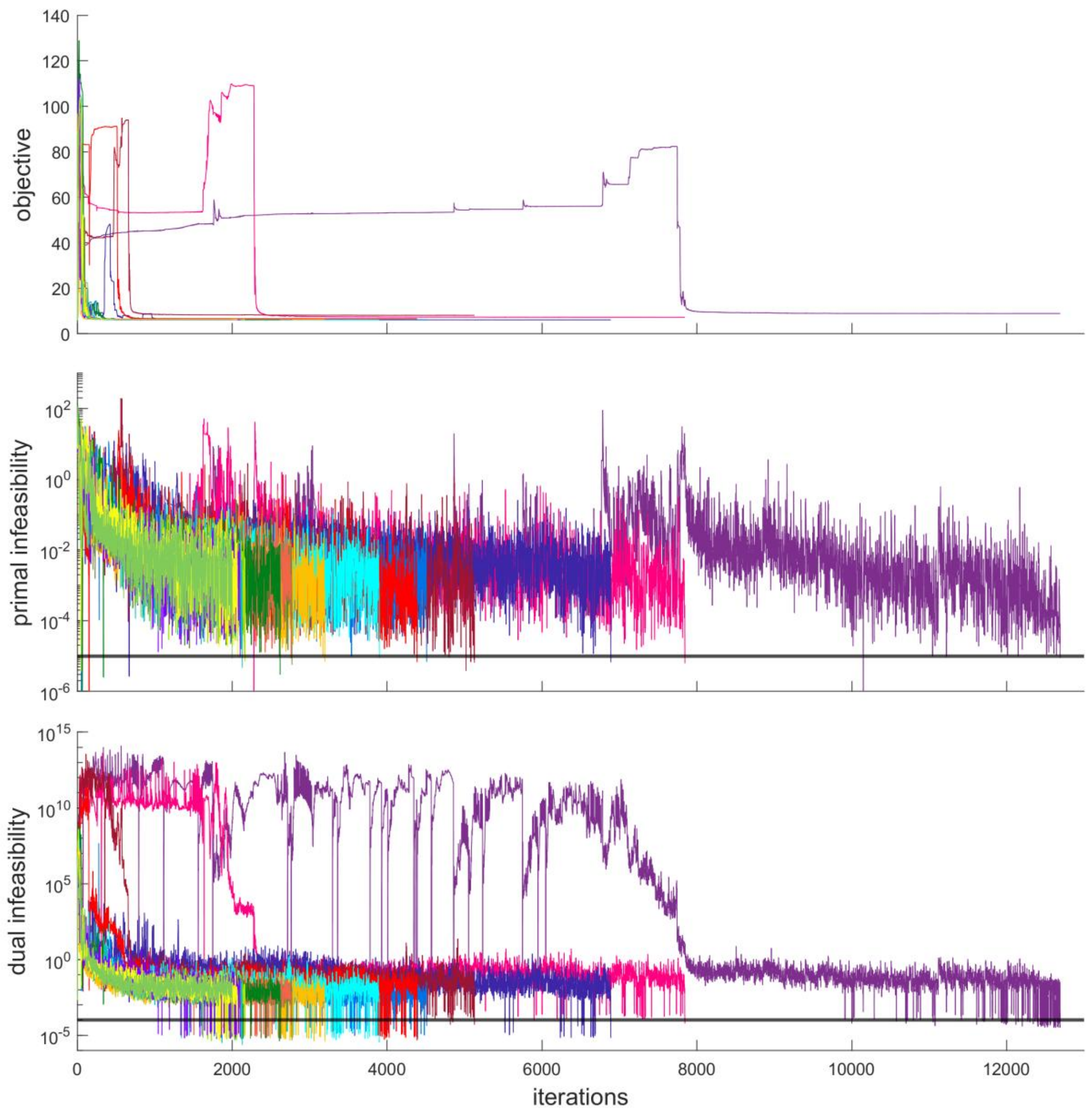


**Fig C. Convergence of the simulations.**
The primal infeasibility is the highest constraint violation, the dual infeasibility is the highest absolute value of the gradient of the Lagrangian (see equation 4 and 5 in (5)). The horizontal black lines indicate the thresholds to accept a solution. When the objective remains similar while the infeasibilities oscillate, this indicates poor convergence towards a solution.

### References


1. Andersson JAE, Gillis J, Horn G, Rawlings JB, Diehl M. CasADi: a software framework for nonlinear optimization and optimal control. Math Program Comput. 2019 Mar 1;11(1):1–36. doi:10.1007/s12532-018-0139-4

2. Lawson CL, Hanson RJ, Kincaid DR, Krogh FT. Basic Linear Algebra Subprograms for Fortran Usage. ACM Trans Math Softw. 1979 Sep 1;5(3):308–23. doi:10.1145/355841.355847

3. AOCL-BLIS [Internet]. AMD; [cited 2025 Nov 25]. Available from: https://www.amd.com/en/developer/aocl/blis.html

4. BLAS (Basic Linear Algebra Subprograms) [Internet]. [cited 2026 Feb 7]. Available from: https://netlib.org/blas/

5. Wächter A, Biegler LT. On the implementation of an interior-point filter line-search algorithm for large-scale nonlinear programming. Math Program. 2006 Mar 1;106(1):25–57. doi:10.1007/s10107-004-0559-y

6. Hogg JD, Scott JA. HSL_MA97 : a bit-compatible multifrontal code for sparse symmetric systems. Rutherford Appleton Lab Tech Rep [Internet]. 2011 [cited 2025 Mar 6];RAL-TR-2011-024. Available from: https://epubs.stfc.ac.uk/work/61445

7. HSL. A collection of Fortran codes for large scale scientific computation. [Internet]. [cited 2026 Feb 8]. Available from: https://www.hsl.rl.ac.uk/
