## Supplementary material for "Stochastic optimal control simulations of walking: potential and perspective": S3 Text

### S3 Text. Additional results

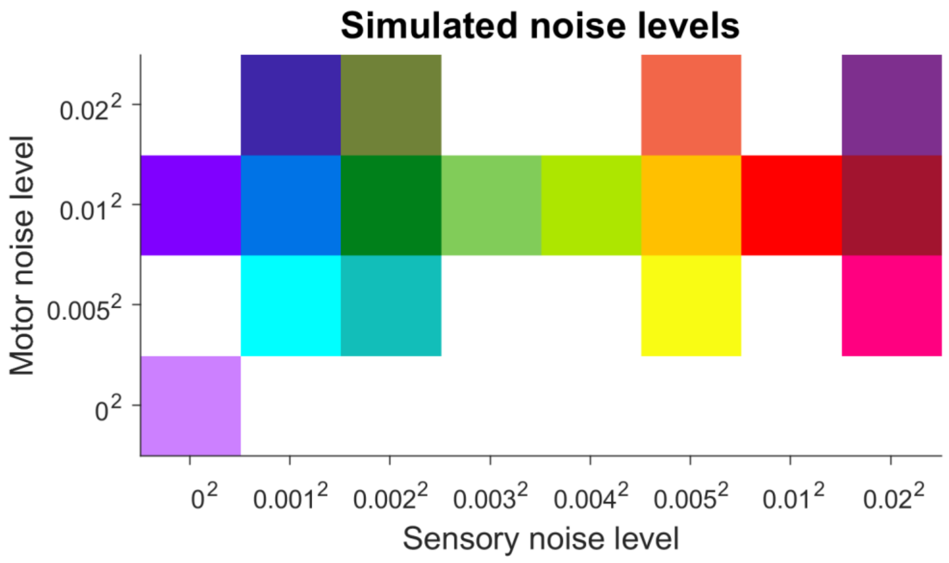


**Fig A. Each simulated combination of sensory and motor noise levels has a specific colour, which is used across figures.**

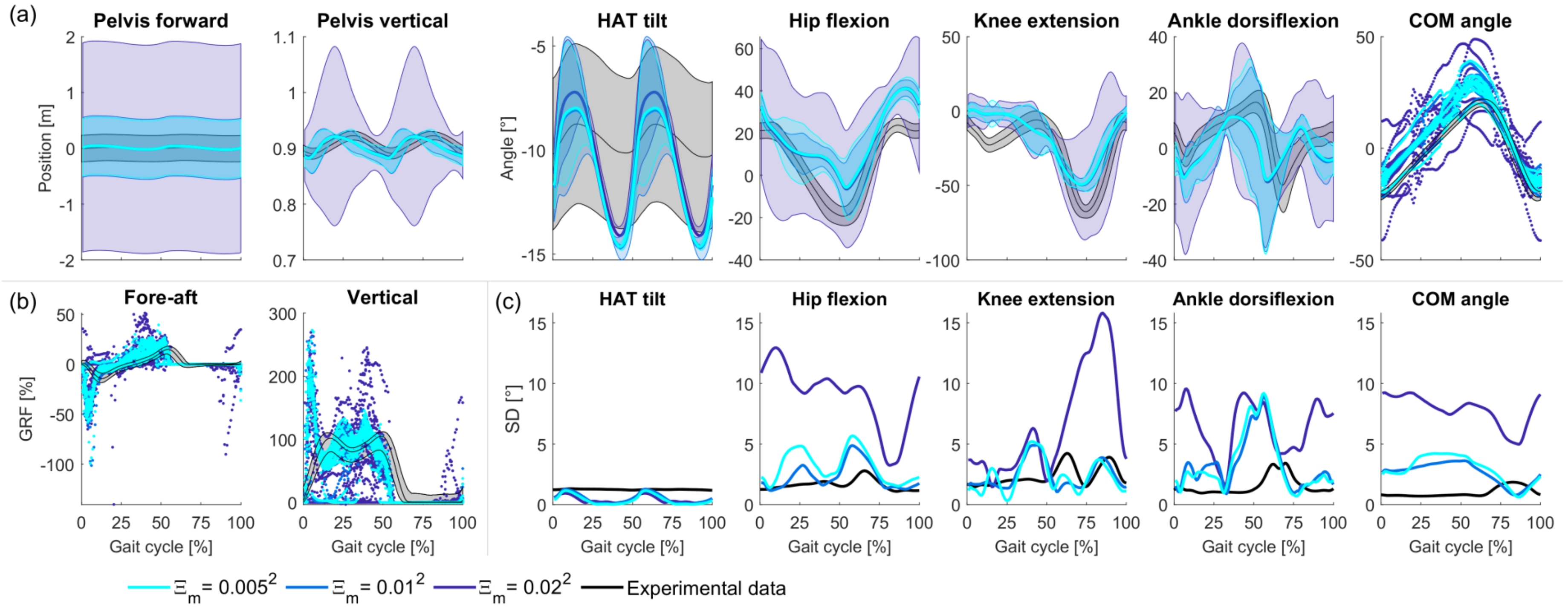


**Fig B. Sensitivity to motor noise level; sensory noise level 0.001$^2$.**
(a) Kinematics. (b) Ground reaction forces. (c) Standard deviation of kinematics.

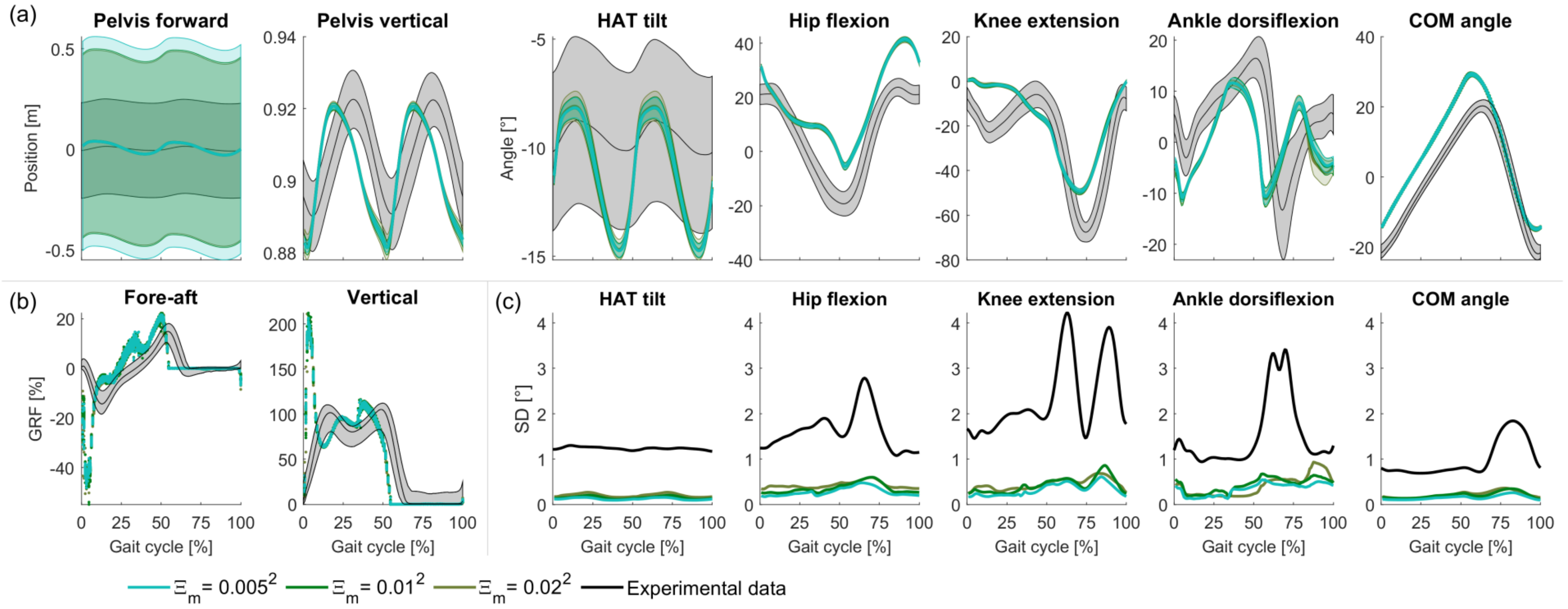


**Fig C. Sensitivity to motor noise level; sensory noise level $0.002^2$.**
(a) Kinematics. (b) Ground reaction forces. (c) Standard deviation of kinematics.

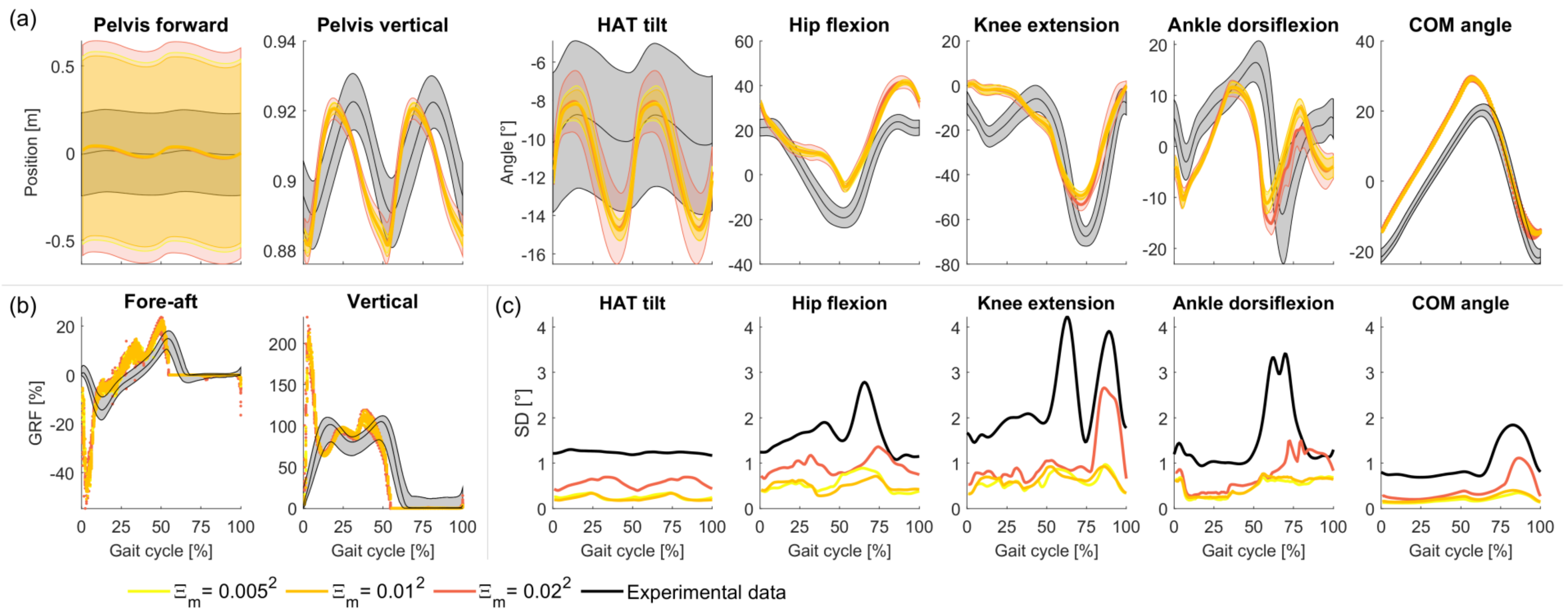


**Fig D. Sensitivity to motor noise level; sensory noise level 0.005$^2$.**
(a) Kinematics. (b) Ground reaction forces. (c) Standard deviation of kinematics.

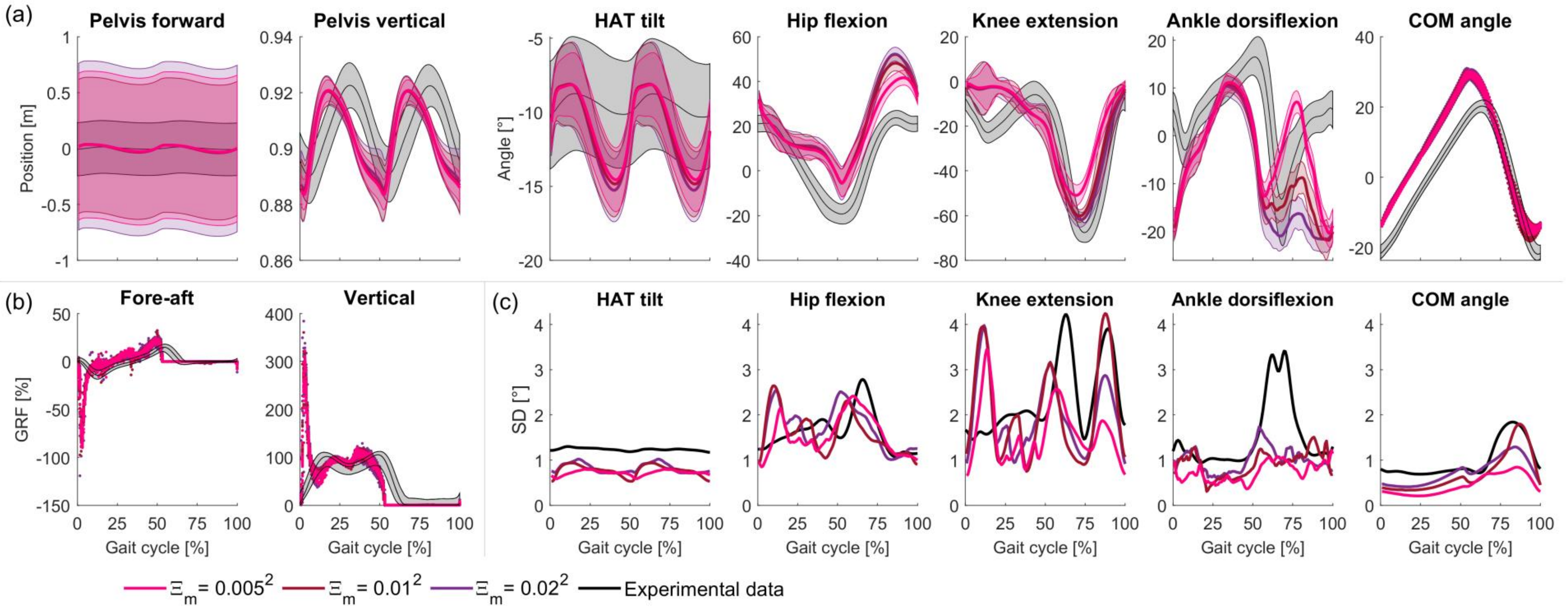


**Fig E. Sensitivity to motor noise level; sensory noise level $0.02^2$.**
(a) Kinematics. (b) Ground reaction forces. (c) Standard deviation of kinematics.

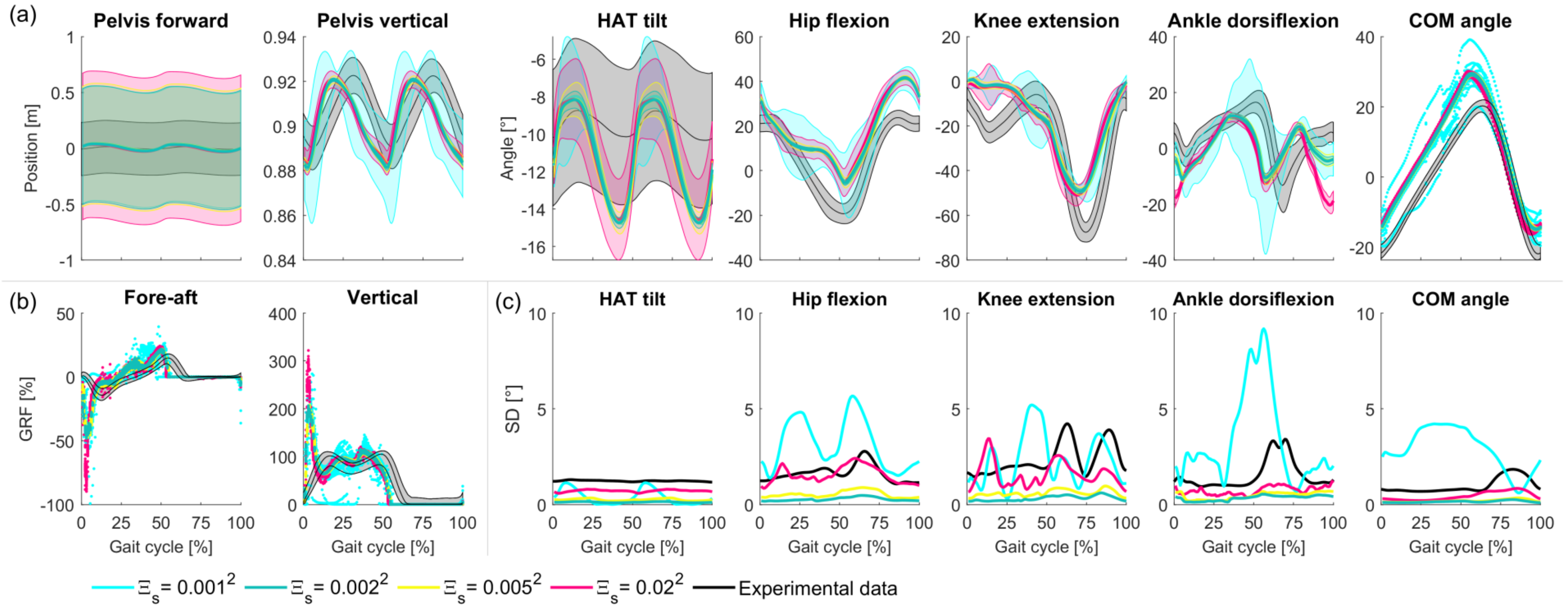


**Fig F. Sensitivity to sensory noise level; motor noise level $0.005^2$.**
(a) Kinematics. (b) Ground reaction forces. (c) Standard deviation of kinematics.

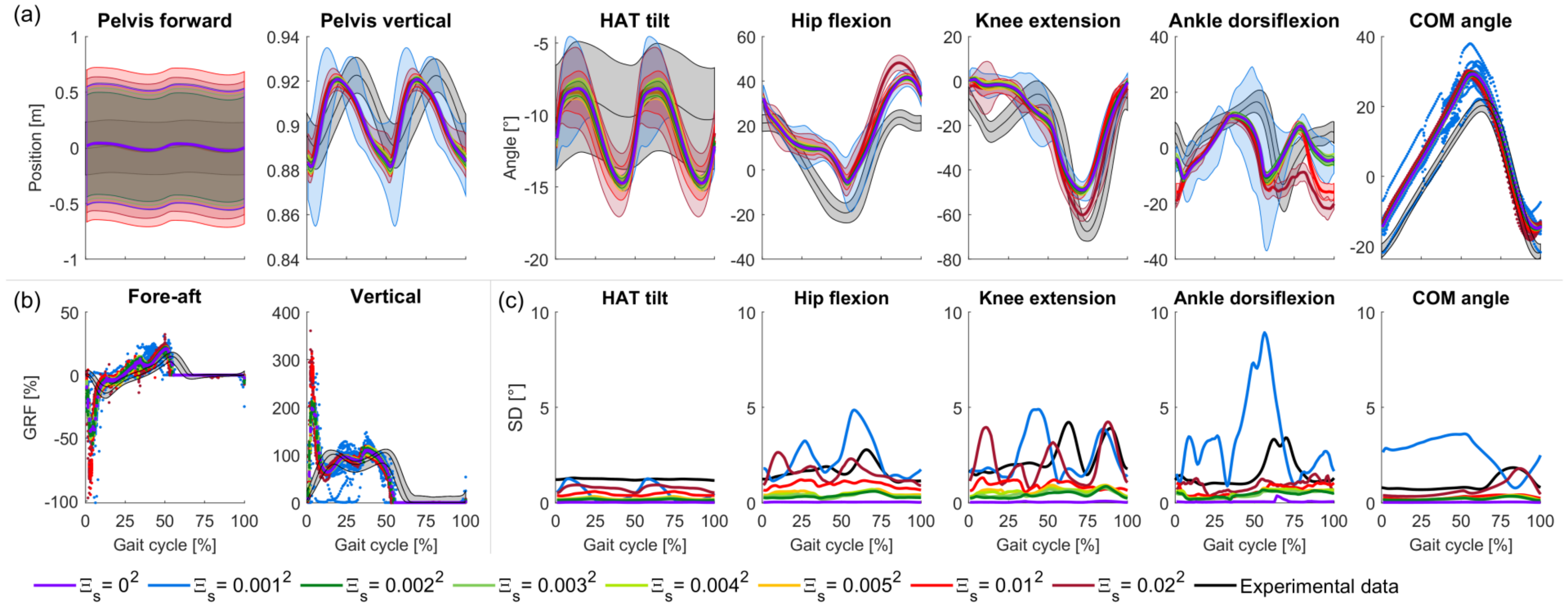


**Fig G. Sensitivity to sensory noise level; motor noise level $0.01^2$.**
(a) Kinematics. (b) Ground reaction forces. (c) Standard deviation of kinematics.

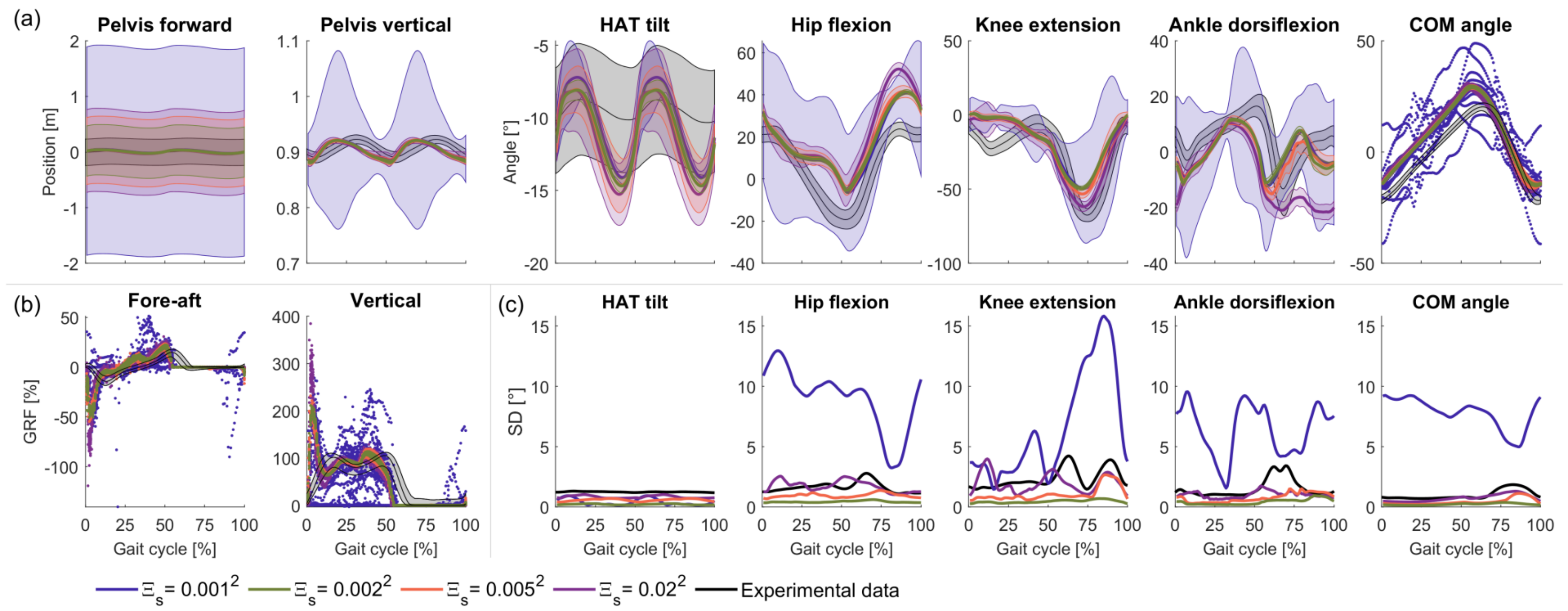


**Fig H. Sensitivity to sensory noise level; motor noise level $0.02^2$.**
(a) Kinematics. (b) Ground reaction forces. (c) Standard deviation of kinematics.

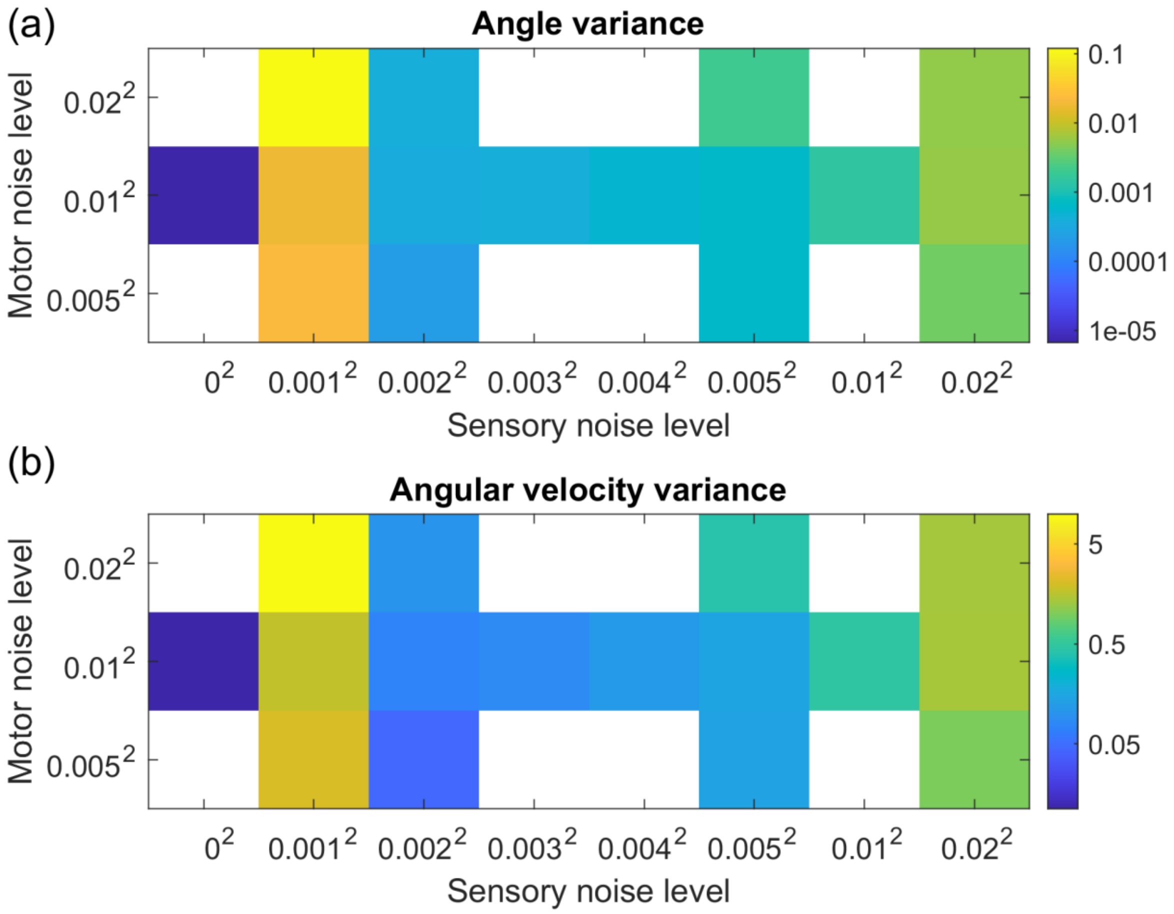


**Fig I. State variance, considering only angles or angular velocities.**

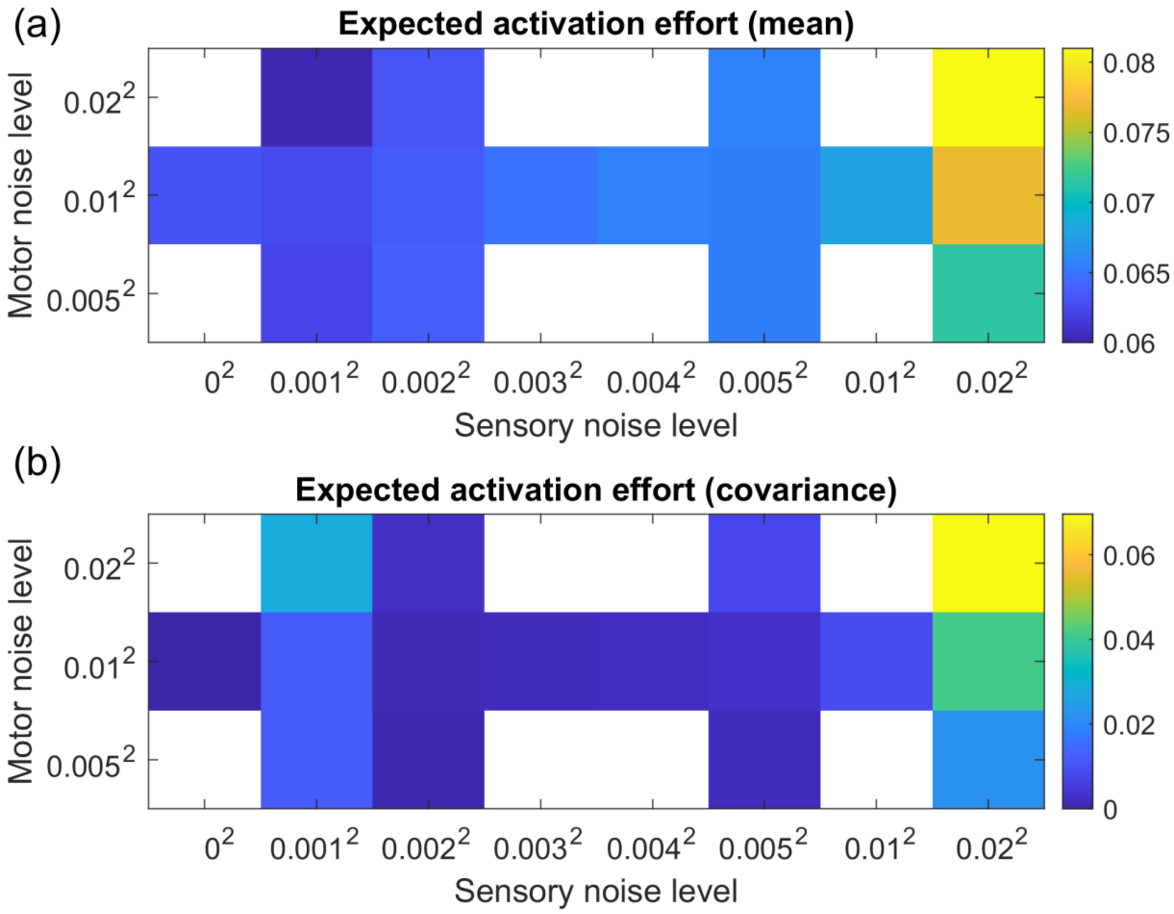


**Fig J. Expected effort due to (a) mean and (b) covariance of muscle activation.**
These are the first two terms of the series expansion of the expected effort (equation 1)

$$\int \mathbb{E}(\|a\|_2^2)\, dt \approx \underbrace{\int \|\bar{a}\|_2^2 dt}_{mean\ term} + \underbrace{\int trace\big(cov(a)\big) dt}_{covariance\ term} + \cdots \tag{1}$$

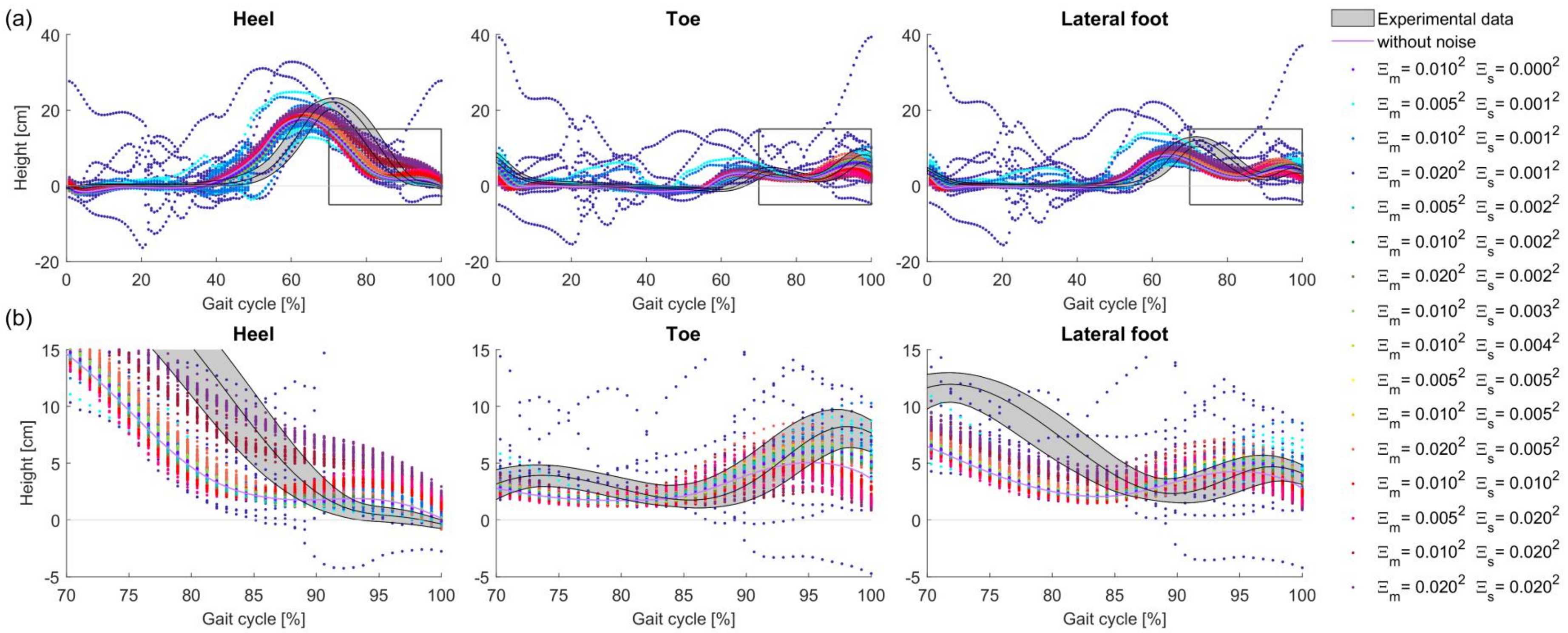


**Fig K. Simulated and experimental foot marker heights.**

(a) Full gait cycle. (b) Detail view of minimal swing foot clearance.

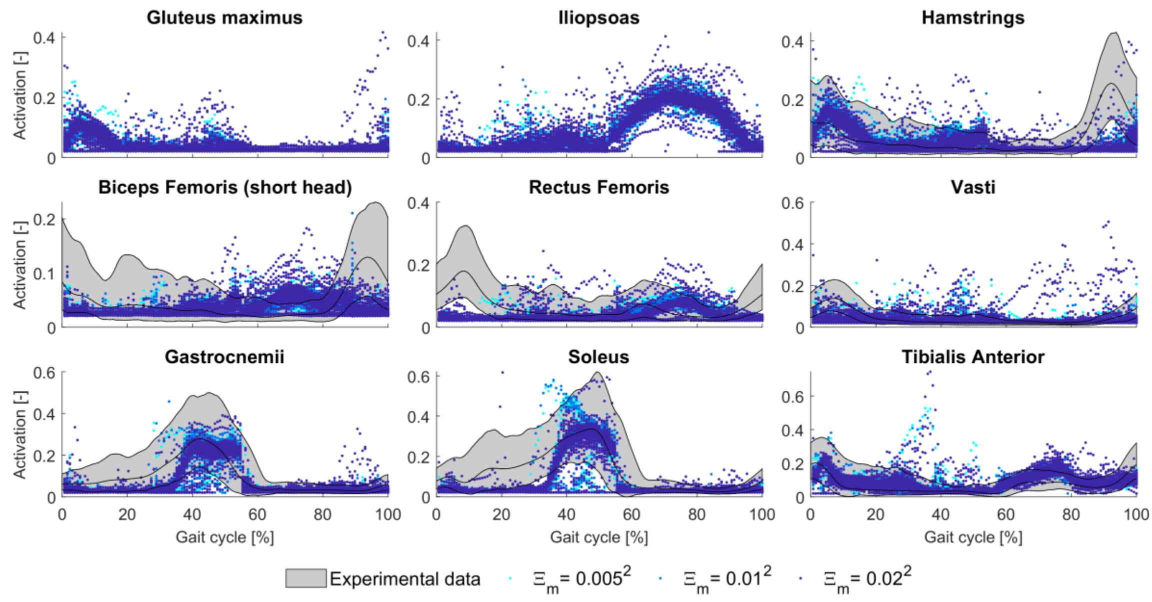


**Fig L. Sensitivity of muscle activations to motor noise level; sensory noise level $0.001^2$.**

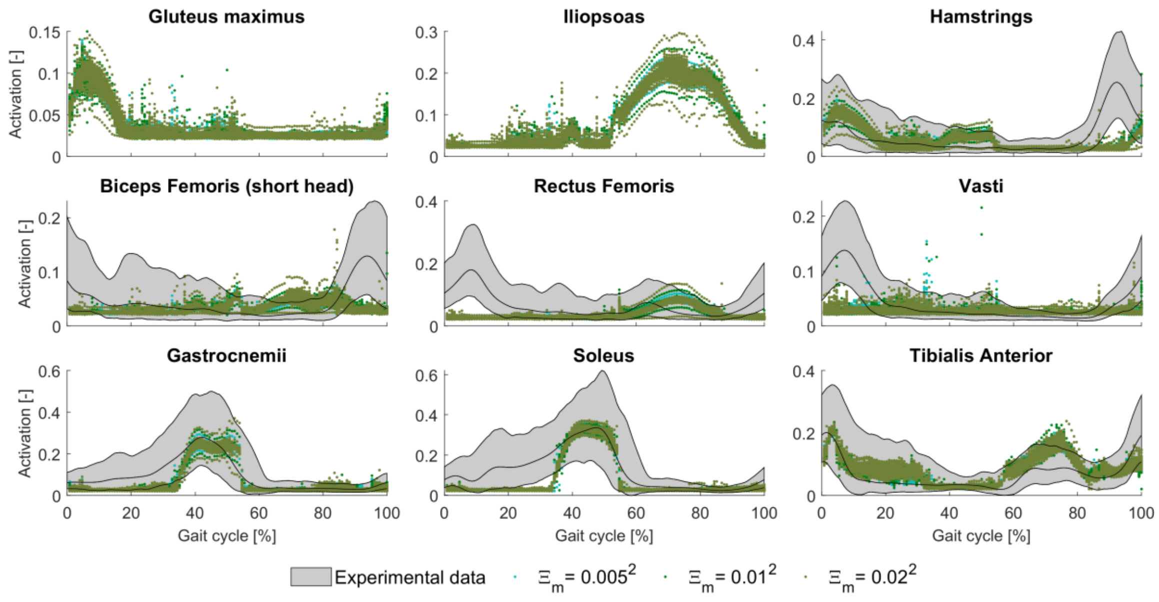


**Fig M. Sensitivity of muscle activations to motor noise level; sensory noise level $0.002^2$.**

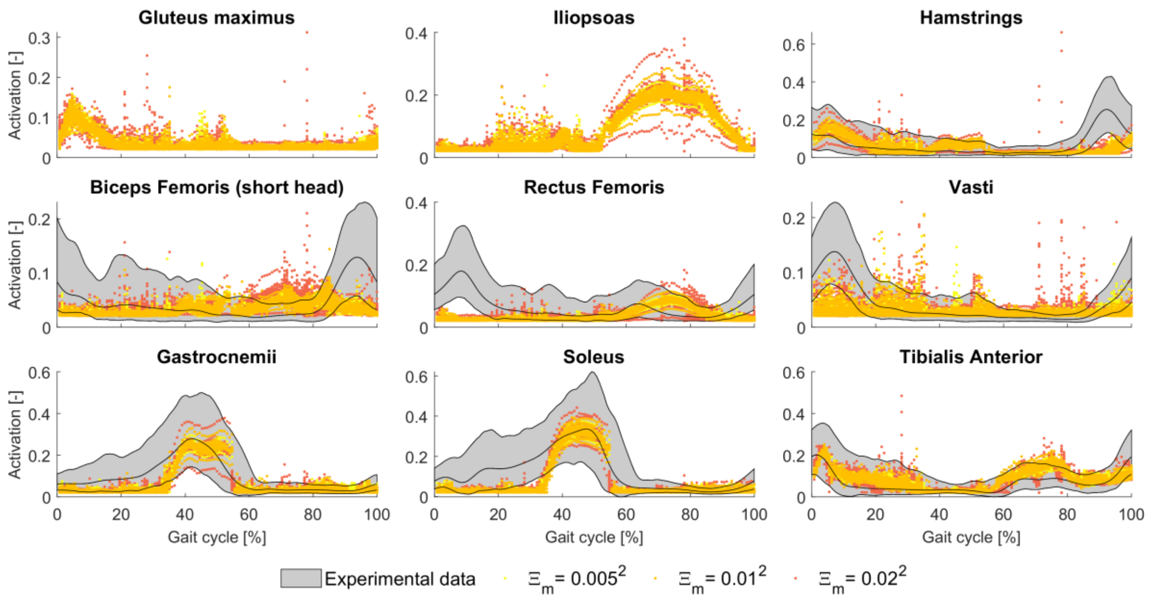


**Fig N. Sensitivity of muscle activations to motor noise level; sensory noise level $0.005^2$.**

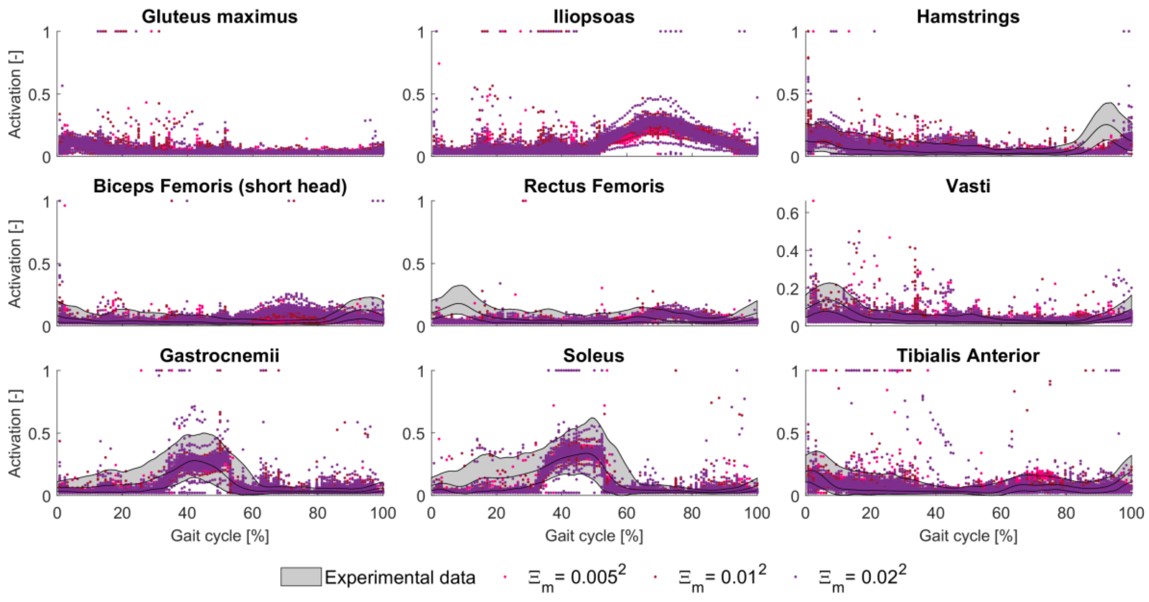


**Fig O. Sensitivity of muscle activations to motor noise level; sensory noise level $0.02^2$.**

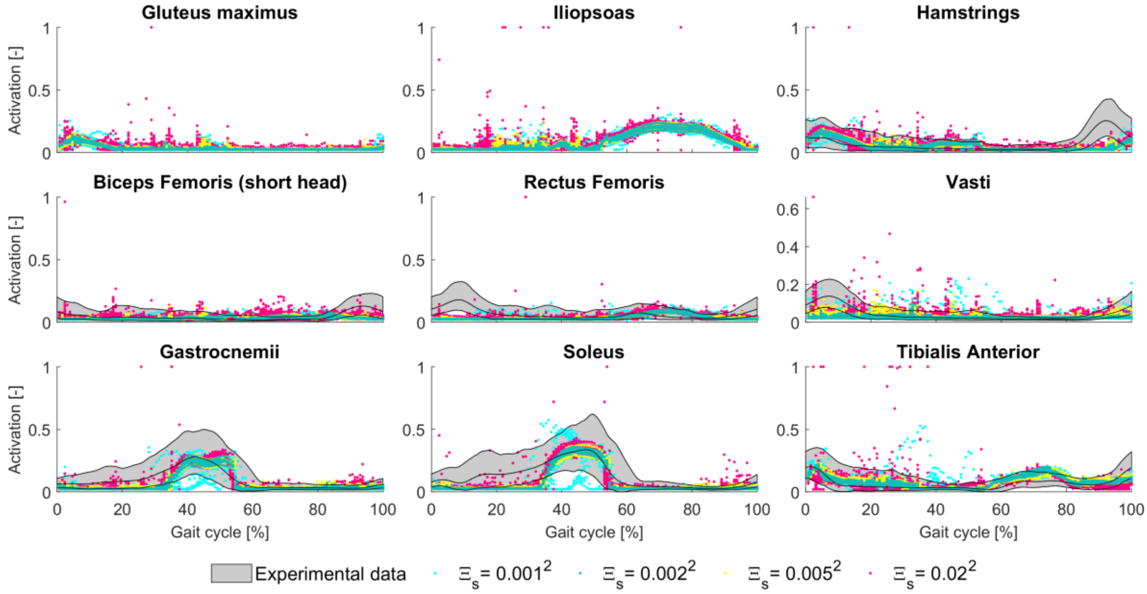


**Fig P . Sensitivity of muscle activations to sensory noise level; motor noise level $0.005^2$.**

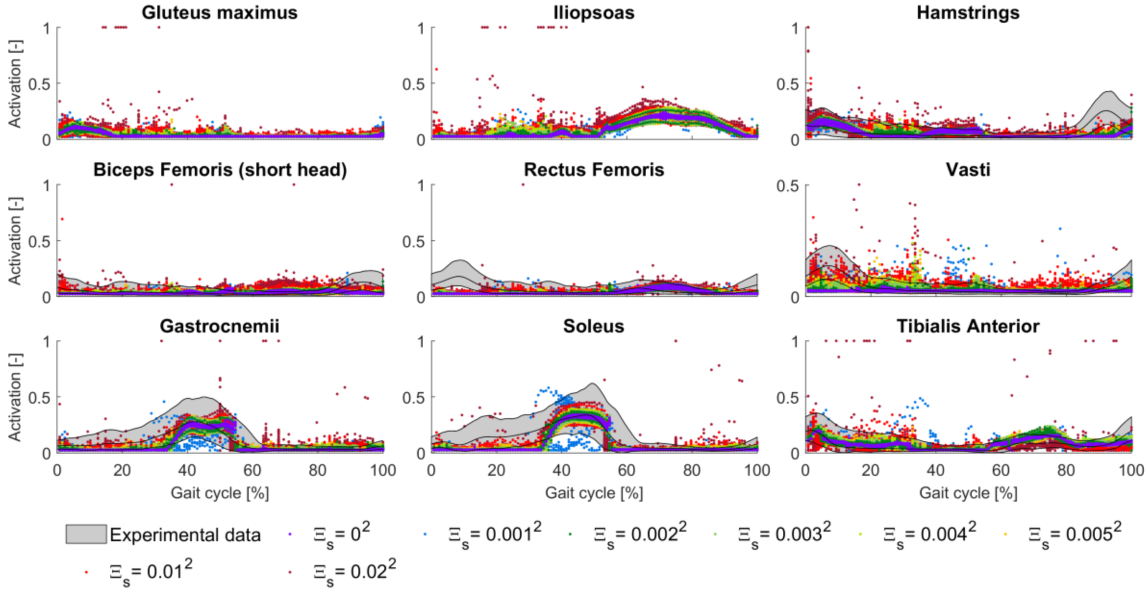


**Fig Q . Sensitivity of muscle activations to sensory noise level; motor noise level $0.01^2$.**

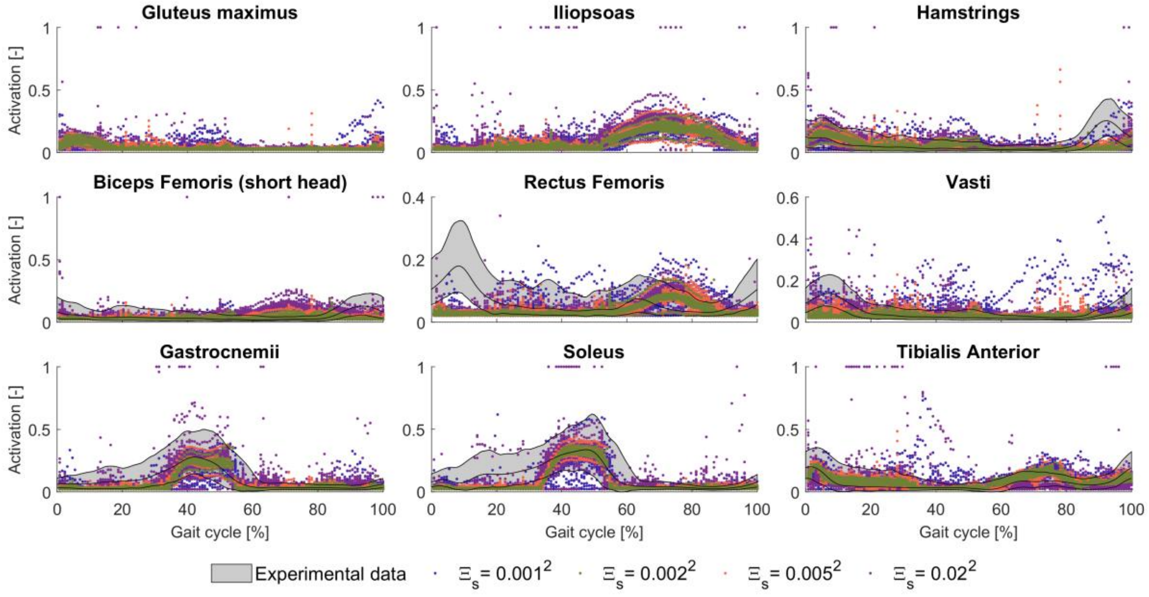


**Fig R . Sensitivity of muscle activations to sensory noise level; motor noise level $0.02^2$.**
